# The ASC Speck and NLRP3 Inflammasome Function Are Spatially and Temporally Distinct

**DOI:** 10.1101/2021.07.19.452947

**Authors:** Abhinit Nagar, Tabassum Rahman, Jonathan A. Harton

## Abstract

Although considered the ternary inflammasome structure, whether the singular, perinuclear NLRP3:ASC speck is synonymous with the NLRP3 inflammasome is unclear. Herein we report that the NLRP3:ASC speck is not required for nigericin-induced inflammasome activation, but facilitates and maximizes IL-1β processing. Further, the NLRP3 agonists H_2_O_2_ and MSU elicited IL-1β maturation without inducing specks. Notably, caspase-1 activity is spatially distinct from the speck, occurring at multiple cytoplasmic sites. Additionally, caspase-1 activity negatively regulates speck frequency and speck size while speck numbers and IL-1β processing are negatively correlated, cyclical, and can be uncoupled by NLRP3 mutations or inhibiting microtubule polymerization. Finally, when specks are present, caspase-1 is likely activated after leaving the speck structure. Thus, the speck is not the NLRP3 inflammasome itself, but is instead a dynamic structure which may amplify the NLRP3 response to weak stimuli by facilitating the formation and release of small NLRP3:ASC complexes which in turn activate caspase-1.

## INTRODUCTION

Inflammasomes are intracellular, multi-protein complexes that assemble and activate capsase-1 following stimuli of microbial, host, and environmental origin (Cheng et al., 2010; Latz et al., 2013). Active caspase-1 cleaves pro-IL-1β and pro-IL-18, cytokines vitally important during infection and inflammation (Lamkanfi and Dixit, 2014; Sharma and Kanneganti, 2016). Caspase-1 also cleaves gasdermin-D (GSDMD) which then forms a membrane pore facilitating IL-1β release and pyroptotic cell-death (Shi et al., 2015). Most inflammasomes comprise members of the NLRP family of intracellular pathogen receptors that associate with the ASC adaptor protein to recruit caspase-1 (Lamkanfi and Dixit, 2014; Latz et al., 2013; Sharma and Kanneganti, 2016) Among NLRP inflammasomes, the NLRP3 inflammasome is essential for innate protection against a plethora of microbial pathogens and contributes to numerous inflammatory pathologies (Lamkanfi and Dixit, 2014; Latz et al., 2013).

ASC-dependent inflammasome activation is accompanied by rapid relocation of the NLR and ASC into a singular, perinuclear, punctate ‘speck’ structure of approximately 1µm (Hoss et al., 2017; Stutz et al., 2013). The speck is thought to act as a supramolecular signaling complex (SMOC), higher-order structures which locally concentrate weakly interacting proteins required for signal transduction (Kagan, 2012; Kagan et al., 2014; Wu, 2013). Speck formation is rapid causing a concomitant drop in cytosolic ASC concentration, approximately 200-fold, to sub-micromolar concentrations within 100 seconds (Cheng et al., 2010) and almost all available NLRP3 and ASC oligomerizes within minutes (Fernandes-Alnemri et al., 2007). Speck assembly is microtubule-dependent (Elliott and Sutterwala, 2015; Li et al., 2017; Misawa et al., 2013). During NLRP3 inflammasome activation, ASC binds to damaged mitochondria and is drawn via microtubules to NLRP3 on the endoplasmic reticulum (Elliott and Sutterwala, 2015). NLRP3 and ASC then associate forming the speck (Bryan et al., 2009; Elliott and Sutterwala, 2015). Accordingly, microtubule depolymerization agents such as colchicine block speck formation (Misawa et al., 2013) and impair NLRP3 inflammasome maturation of IL-1β induced by monosodium urate crystals (Martinon et al., 2006). Thus, colchicine is a drug of choice for treating gout, an NLRP3-related inflammatory condition (Dalbeth et al., 2014).

Since NLRP inflammasome activity and speck assembly are concomitant and both require interaction between the NLRP and ASC, NLRP3:ASC specks are generally equated with the inflammasome (Elliott and Sutterwala, 2015). Studies using small molecules including colchicine, nocodazole, and others which prevent both speck assembly and inflammasome function support this view (Coll et al., 2015; Misawa et al., 2013). However, biochemical analysis suggest active NLRP1:ASC:caspase-1 complexes may be less than ∼0.45 μm in diameter (∼700 kDa) (Martinon et al., 2002), values generally accepted for NLRP3 complexes as well (Lu et al., 2014) and are much smaller than the 1µm speck structure. Likewise, in vitro assembled NLRP1 (15 Å) and NLRP3 (100 nm) inflammasome structures have an approximately 100-1000-fold smaller diameter than the speck, different arrangements, and distinct stoichiometries (Faustin et al., 2007; Lu et al., 2014). Further, while NLRP3:ASC speck formation requires a few minutes, IL-1β processing is detected within 1 hour (Martinon et al., 2004). Such size, stoichiometric, and temporal disparities between the speck and the inflammasome suggest these structures may have distinct functions. Nevertheless, the relationship between the inflammasome and the speck is still unclear. Whether such distinctions reflect separate functions for the speck and inflammasome is unanswered.

This study establishes that NLRP3-mediated caspase-1 activity (the inflammasome) is distinct and independent of the NLRP3:ASC speck. While not required and insufficient for NLRP3 inflammasome function, the NLRP3:ASC speck is a dynamic, spatially and temporally distinct structure regulated in part by caspase-1. Further, the speck lowers the NLRP3 inflammasome activation threshold, potentially amplifying the formation and release of much smaller inflammasome complexes. The implications of our data and the relationship between ASC speck structures and the NLRP3 inflammasome are discussed.

## MATERIALS AND METHODS

### Cell Culture

Human kidney epithelial cells (HEK239T) (ATCC; Cat.# CRL-11268) and immortalized bone-marrow derived macrophages (iBMDMs) (a gift from Dr. Katherine Fitzgerald, UMass Medical School, Worcester, MA) were cultured in complete Dulbecco’s Modified Eagle Medium/high glucose (DMEM) (Hyclone™; Cat.# SH30022.01) supplemented 10% FBS (Atlanta Biologicals; Cat.# S11050), 1X Glutamax (Gibco; Cat.# 35050-061), and 0.1% penicillin/Streptomycin at 37°C, 5% CO_2_. Primary human monocytes were obtained from University of Nebraska Medical Center and cultured in DMEM supplemented with 10% Human AB Serum (Corning; Cat.# 35-060-Cl). THP-1 (ATCC; TIB-202) monocytic cells were cultured in RPMI-1640 (Hyclone™; Cat.# SH30027.01) supplemented with 10% FBS, 1X-β-mercaptoethanol (Gibco; Cat.# 21985) and 1X-Glutamax. THP-1 cells were differentiated to macrophages by treatment with PMA (100 nM) for 72 hrs. Cultured cell lines were maintained at sub-confluence and split every 2-3 days. Cell numbers and viability were determined by trypan blue exclusion.

### Expression plasmids, construct and DNA transfection

Expression plasmids encoding NLRP3, NLRP3 cysteine mutants, and chimeric NLRP3 mutants, myc-ASC, GFP-ASC, caspase-1, and pro-IL-1β have been described previously (Atianand et al., 2011; Nagar et al., 2019; Rahman et al., 2020). GFP-YVAD was PCR-amplified from pEGFP-C3 and cloned into the Lamp1-RFP plasmid (a gift from Walter Mothes; RRID: Addgene Cat.#1817) (Sherer et al., 2003) as an EcoRI (New England Biolabs (NEB); Cat.# R0101S)/BamHI (NEB; Cat.# R0136S) fragment to generate GFP-YVAD-RFP **(Fig. S1)**. Ligation was performed using T4 DNA Ligase (Promega; Cat.# M1801) using manufacturer’s protocol for staggered end ligation. Caspase-1 mutants **(Fig. S2A)** were generated by QuikChange PCR. The parental caspase-1 plasmid (methylated) was digested using DpnI (NEB; Cat,# R0176S). Plasmids were transformed in *E*.*coli* DH5α competent cells (NEB; Cat,# C2987I). All site-directed changes were confirmed by sequencing. Oligonucleotide primers used in this project were produced by Integrated DNA Technology and are listed in **Table 1**. DNA transfections were carried out using FuGENE6 (2.5 μl/μg DNA) (Roche Applied Science; Cat.# 11988387001) as per manufacturer’s protocol.

**Table 1:**
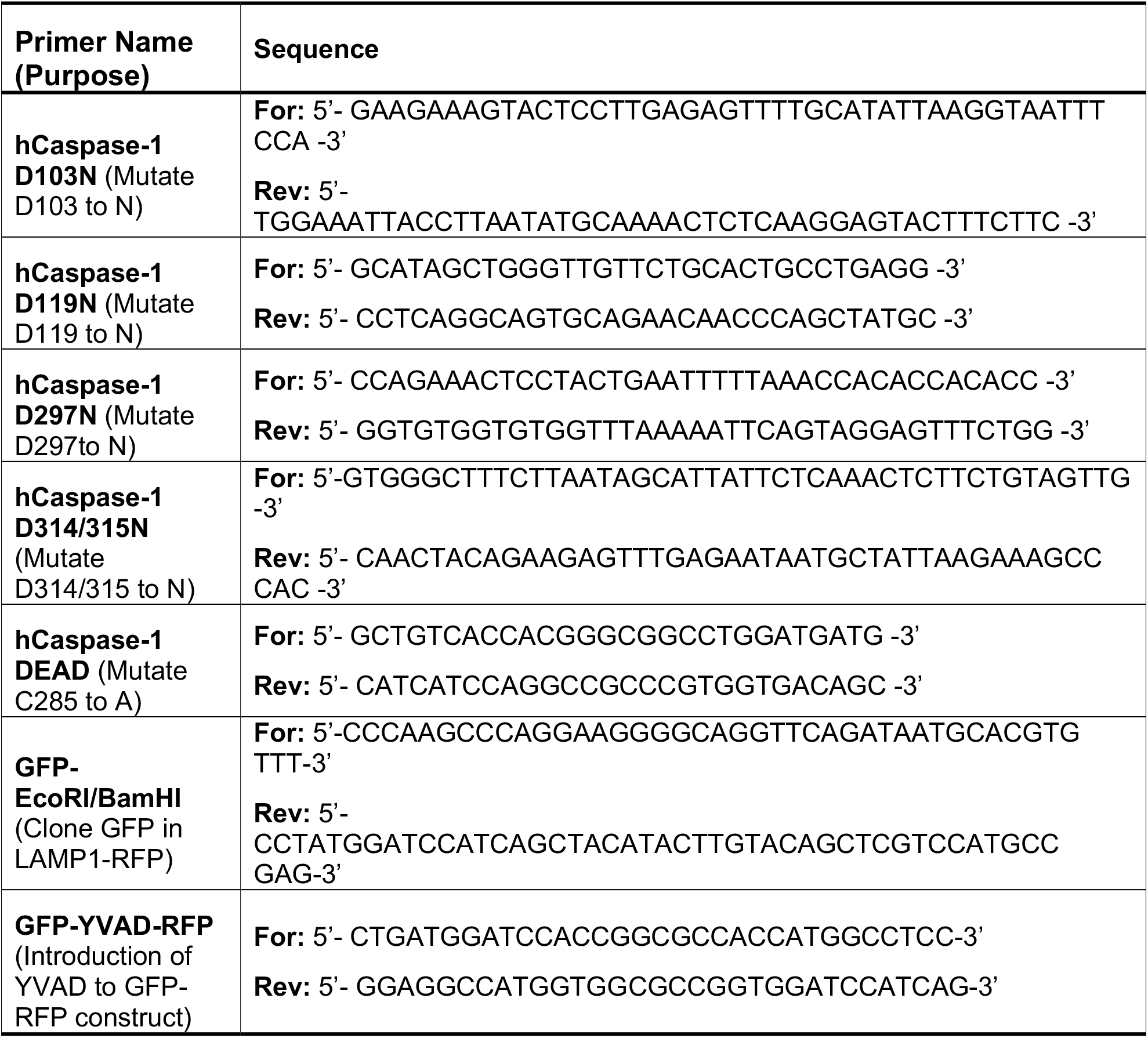
Primers used in the study.

### Transfection conditions

For assays of stimulus-dependent NLRP3-inflammasome IL-1β maturation and release (inflammasome reconstitution), HEK 293T cells were seeded (2.5 × 10^4^) cells/well/ml in 24-well plates, cultured overnight, and transfected with 40 ng pro-caspase1, 200 ng pro-IL1β, 8 ng ASC, and 100 ng of either an NLR or control (pcDNA3) plasmid. Under these conditions, specks are not evident and NLRP3 and ASC cannot be detected by immunofluorescence staining due to low levels of expression.

Observing macrophage-like, agonist-dependent speck formation in HEK293T cells either alone (using Time-of-Flight Inflammasome Evaluation (TOFIE)) or in conjunction with caspase-1 activity (using Inflammasome and Caspase-1 activity Characterization and Evaluation (ICCE)) requires higher levels of expressed NLRP3 and ASC. 2×10^5^ HEK293Ts were seeded per well/ml in 12-well plates and after overnight incubation, transfected with 100 ng NLRP3, 50 ng GFP-ASC, either alone or with caspase-1 (20 or 50 ng). Empty vector, pcDNA3, was used to adjust the final amount of DNA to 1 μg prior to transfection.

For evaluation of speck formation capacity by microscopy, 5×10^5^ HEK293Ts were seeded per well/ml in 6 well plates with cover-slips, after overnight incubation, transfected with 1 µg NLRP3 or its chimeric mutants, and 1 µg ASC and incubated for 20 hrs.

### FRET and Caspase-1 activation assay

The specificity of caspase-1 is significantly increased if peptide flanking the caspase-1 specific sequence YVAD is increased by only four amino acid sequences (Thornberry et al., 1992). Thus, a double reporter (GFP-YVAD-RFP) construct with the sequence “YVAD” separating GFP and RFP was generated, which is expected to have higher specificity than FLICA **(Fig. S1)**. 2×10^5^ HEK293Ts were seeded per well/ml in 12 well plates, after overnight incubation, transfected with 100 ng NLRP3, 100 ng GFP-ASC, 400 ng p-Casp-1 (WT or mutant) and GFP-YVAD-RFP. As a positive control for FRET, cells were transfected with all the above stated plasmids except capase-1 and for negative control, cells were transfected with GFP and RFP expressed on different plasmids instead GFP-YVAD-RFP expressing plasmid. Transfections were carried out at a constant DNA concentration of 1μg/well. Empty vector, pcDNA3, was used to adjust the final amount of DNA to 1μg. After overnight incubation, cells were either analyzed by microscopy to evaluate the distribution of GFP-YVAD-RFP reporter. To measure caspase-1 activation using FRET, cells were harvested after overnight incubation and fixed using 4% PFA as described above. The cells were acquired on LSRII flow cytometer equipped with 405 nm, 488 nm, and 642 nm lasers with long-pass filter of 505 nm and band-pass filters of 450/50 nm, 530/30 nm, and 660/20 nm. Acquisition was done using BD FACSDiva software. Data was analyzed using FlowJo. Samples were gated to exclude debris and cell doublets. Singlet population was further gated for GFP staining. A stop gate of 10^4^ cells was set on the GFP-positive gate. GFP+ cells were plotted for RFP vs GFP staining. Since the flow cytometer lacked lasers to excite RFP, only source for RFP signal was through GFP emission leading to FRET. Thus, events in FRET channel was normalized to 100 for GFP-YVAD-RFP transfected sample and normalized to 0 for GFP+RFP transfected samples. FRET was calculated using the following formula:

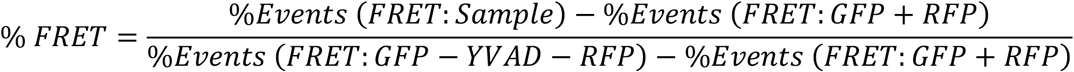

For caspase-1 activity measurements, caspase-1 transfected sample was normalized to 100 and GFP-YVAD-RFP alone transfected sample was normalized to 0. Caspase-1 activity was measured using the following formula:

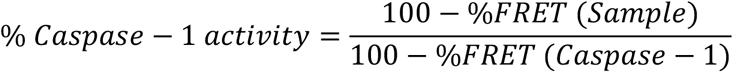

### Inflammasome activation

Inflammasome reconstituted HEK293T cells were infected with *Francisella novicida* U112 (Albany Medical College Microbiology Core Facility) at an moi of 100, 4 hrs post-transfection. After 24 hrs, culture supernatants were collected by centrifugation. For H_2_O_2_ (Sigma Aldrich; Cat.# 216713), nigericin (Invivogen; Cat.# tlrl-nig), and monosodium urate (MSU) (Sigma Aldrich; Cat#U-2875) stimulation, reconstituted HEK293T cells were treated 18 hrs post-transfection with 100 µM H_2_O_2_ for 1 hr, with 5 µM nigericin for 2hrs or with 150 µg/ml MSU for 2 hrs and culture supernatants were collected. Control untreated wells were harvested at the same time. Secreted IL-1β was measured with the Human IL-1β cytoset ELISA (Invitrogen; Cat.# CHC1213) as per the manufacturer’s instructions. THP-1 cells, immortalized BMDMs, and primary human monocytes were treated with 100 ng/ml LPS (O26:B6) (Sigma Aldrich; Cat.# L2654) for 3-4 hrs. Following LPS treatment cells were stimulated with 5 mM ATP (Sigma Aldrich; Cat.# A3377) or 10 µm nigericin for 30 mins or 150 µg/ml MSU for 2 hrs. For microscopy, THP-1 cells were treated with 100 nM PMA for 72 hrs, washed, and then supplemented with 500µl serum-free RPMI media containing vehicle or 5 µm colchicine. After 1hr, cells were stimulated with 500µl serum-free RPMI media containing nigericin.

### Time of Flight Inflammasome Evaluation (TOFIE)

A portion of the samples prepared for ISXII analysis (below) was diluted to 200μl in ISXII cell suspension buffer and acquired on an LSRII flow cytometer equipped with 405nm, 488nm, and 642nm lasers with 505nm long-pass filter of and band-pass filters of 450/50nm, 530/30nm, and 660/20nm. BD DIVAS software was used for acquisition and data was analyzed using FlowJo. Samples were gated to exclude debris and cell doublets. The single cell population was further gated for GFP staining. A stop gate of 10^4^ cells was set on the GFP-positive gate. The percentage of cells containing an ASC speck was determined by analyzing the height (H), width (W), and area (A) of the GFP pulse area (high H:A and low W:A indicates speck positive cells) as described previously by Sester and group (Sester et al., 2015; Sester et al., 2016).

### Inflammasome and Caspase-1 activity Characterization and Evaluation (ICCE)

The protocol for ICCE is described in our previous study (Nagar et al., 2019). Following inflammasome activation, media from each well was aspirated leaving 150 μl and volume was made up to 200 μl with media containing cell permeable FLICA660-YVAD-FMK (FLICA660) (1:45 final dilution) (Immunochemistry Technologies; Cat. #9122) and incubated for 30 mins at 37°C with 5% CO_2_. Cells were washed twice with 1X wash buffer (supplied with the FLICA kit). For ImageStream®^X^ Mark II (ISXII) acquisition, cells were treated with 50 µl trypsin-EDTA (Corning; Cat.#25-053-Cl) per well and fixed with 4% paraformaldehyde (PFA) (EMS; Cat.#15710) for 15 mins at room temperature. Following inflammasome activation and FLICA-600 treatment, THP-1 cells, immortalized wild-type and caspase-1/11^-/-^ and primary human monocytes were fixed with 4% paraformaldehyde (PFA) for 15 mins at room temperature, permeabilized with 0.1% TritonX-100 (Fisher Scientific; Cat.# BP151-100) for 10 mins at room temperature and blocked in PBS containing 5% fish gelatin (Sigma Aldrich; Cat.# G7765), 1% BSA (Fisher Scientific; Cat.# BP1605-100), and 0.05% Triton X-100 for 1hr at room temperature. After blocking, cells were stained with Rabbit anti-ASC (N15)-R (1:250) (Santa Cruz; Cat.# sc-22514-R) in wash buffer (PBS containing 1% fish gelatin, 1% BSA, and 0.5% Triton X-100) for 2 hrs. Cells were washed 3 times with wash buffer, followed by incubation with Alexa Fluor®488 goat-anti-rabbit IgG (1:500) (Life Technologies; Cat.# A11034) or Alexa Fluor®594 goat-anti-rabbit IgG_2a_ (1:1000) (Life Technologies; Cat.# A21135) in wash buffer for 1 hr. Fixed cells were harvested and stained with 1 μg/ml DAPI (Invitrogen; Cat.# D-1306) in 1X PBS supplemented with 0.5 mM EDTA (cell suspension buffer) for 10 mins at room temperature (RT). Cells were washed once with 1X PBS and resuspended in 50 μl ISXII cell suspension buffer by gently tapping the tubes (use of a pipette or vortex mixer should be avoided at this stage). The samples were then acquired on the AMNIS Imagestream ISXII.

### Colocalization

Co-localization of proteins with a resolution of 0.5 μm per pixel in the X and Y axis can be determined with the AMNIS ISXII using Bright Detail Similarity (BDS)-index scores, a log-transformed Pearson’s correlation coefficient (Beum et al., 2006). A BDS-index score ≥1.5 is considered significant for colocalization (Wortzel et al., 2015). BDS index was calculated using the ASC speck mask (Mask 14) and FLICA spot mask (Mask V) on a pixel by-pixel and cell-by-cell basis with the Amnis IDEAS Co-localization Wizard tool. Using events from population S as an internal negative control, a threshold BDS-index of 1.5 was set to determine colocalization of speck and active caspase-1 (FLICA) aggregates.

### Microscopy

For microscopy, THP-1, cells were fixed and stained as described for ICCE. NLRP3 was stained using Cryopyrin (H-66) rabbit polyclonal IgG (1:1000) (Santa Cruz; Cat.# sc-66846) or Mouse Monoclonal ANTI-FLAG^®^ M2 antibody (1:1000) (Sigma Aldrich; Cat.# F1804) while performing experiments in HEK293Ts. HEK293T cells were fixed with 4% PFA and washed 3 times with 1X PBS, dipped in distilled water, and mounted on the slide using 10 μl of FLUORO-GEL II mounting medium with DAPI (EMS; Cat.# 17985-51) and visualized using an Axio Observer Z1 fluorescence microscope (Zeiss). Images were acquired using ZEN-Blue at an optimal setting to avoid saturation. Image acquisition was run at identical setting. Image analysis was performed using Fiji-ImageJ software. ASC-positive cells containing specks were counted manually from randomly selected fields acquired at 20X or 60X magnifications. The percentage of speck containing cells was calculated as the fraction of ASC-positive cells containing specks. The percentage of cells containing GFP-RFP aggregates was calculated as the fraction of GFP-RFP-positive cells containing none, single or multiple aggregates.

### Quantification and Statistical Analysis

At least three independent experiments were performed with two/three technical repeats unless otherwise indicated. Statistical test used for data analysis are stated in the figure legends. A p-value of ≤0.05, was considered statistically significant. All statistical analyses were performed using Prism 7.

## RESULTS

### Colchicine prevents speck formation, but not inflammasome activation

Treatment of macrophages with nigericin concentrations above 3 µM stimulates maturation and release of IL-1β even in the presence of colchicine, a microtubule inhibitor that blocks speck assembly (Gao et al., 2016). Therefore, we considered that higher concentrations of nigericin might activate the inflammasome in the absence of the speck, but whether colchicine blockade of NLRP3:ASC speck assembly is overcome under these conditions is untested. PMA-treated THP-1 macrophages were stimulated with increasing concentrations of nigericin in the presence or absence of colchicine. In the absence of colchicine, nigericin stimulation led to maximal IL-1β processing (∼30-fold over untreated controls) irrespective of the dose of nigericin **(Fig. 1A)**. At lower concentrations of nigericin (1 and 2.5 µM), IL-1β maturation and release was blocked by colchicine treatment, but 5 μM nigericin overcame this blockade **(Fig. 1A)**. Although somewhat reduced, the IL-1β response to 5 µM nigericin in the presence of colchicine was approximately 20-fold higher than the untreated control and 65% of that seen without colchicine. While speck formation was expected to accompany inflammasome activity, less than 1% of colchicine-treated cells had ASC specks despite stimulation with 5 µM nigericin **(Fig. 1B&C)**. Thus, IL-1β processing can occur in the absence of specks and speck formation may not be essential for the NLRP3 inflammasome response.

**Figure 1:**
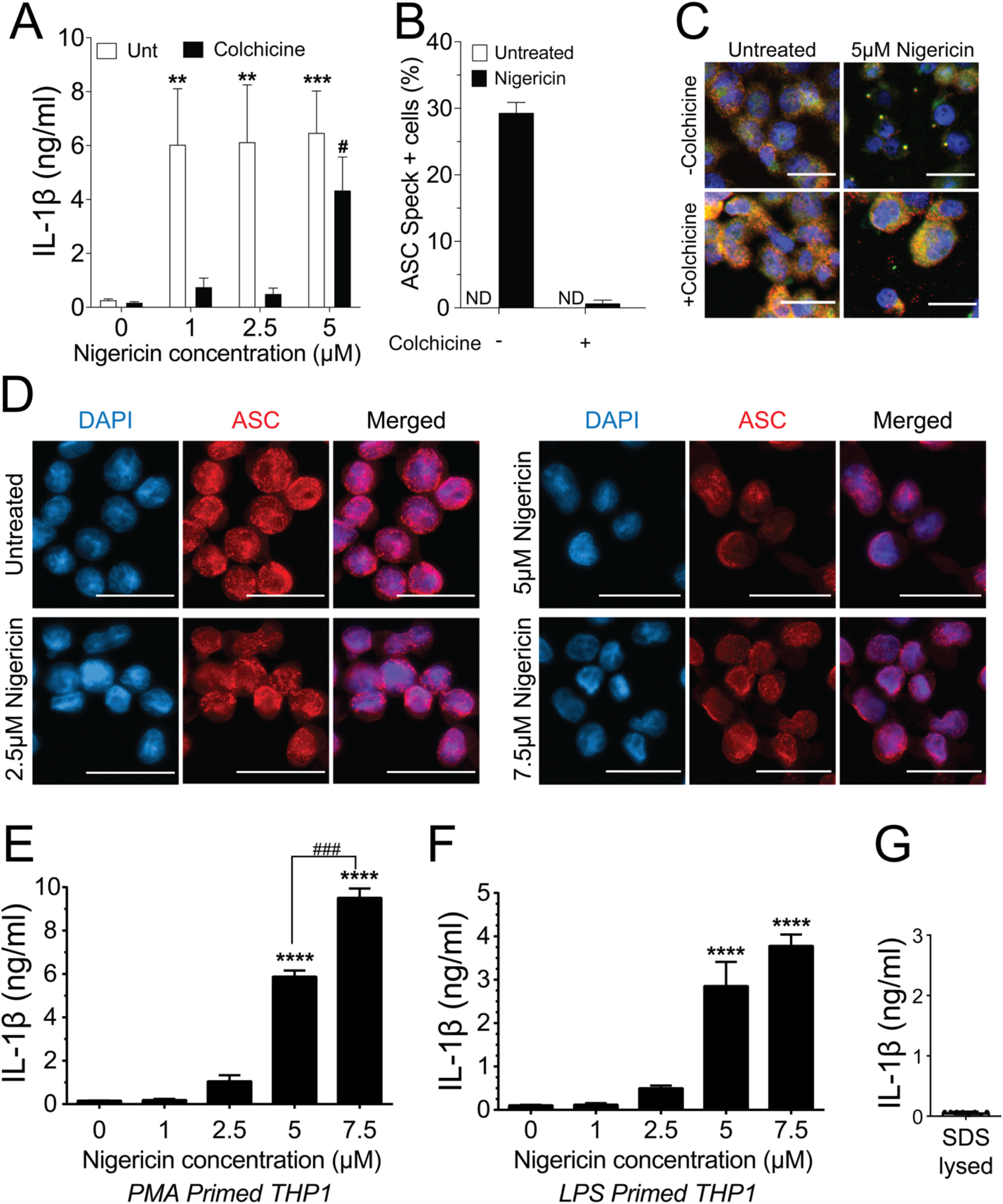
Higher doses of nigericin induce IL-1β release in the absence of specks. **A**. THP-1 cells were treated with 100 nM PMA for 72 hrs and treated with 5 μM colchicine for 1 hr followed by stimulation with different dose of nigericin for 2 hrs. Culture supernatant was used for IL-1β ELISA. **B**. THP-1 cells were treated as described in A. Following nigericin stimulation, cells were fixed and stained for NLRP3 and ASC and analyzed for specks by microscopy. A minimum of 600 cells were analyzed per condition and counted for presence of speck manually. Data represented as mean ± SEM for two independent experiments. N.D.-Not detected. **C**. Representatives micrographs for data shown in B. Cells were stained for NLRP3 (red), ASC (green) and DAPI (blue). Scale bar = 25 nm. **D**. Representatives micrographs for data shown in E. Cells were stained for NLRP3 (red), ASC (green) and DAPI (blue). Scale bar = 25nm. **E**. THP-1 cells were primed with 100 nM PMA for 72 hrs and treated with 5 μM colchicine for 1 hr followed by stimulation with nigericin at the indicated concentrations for 2 hrs. Culture supernatant was used for IL-1β ELISA. **F**. THP-1 cells were treated with 100 ng/ml of LPS for 3-4 hrs and treated with 5 μM colchicine for 1 hrs followed by stimulation with nigericin at the indicated concentrations for 2 hrs. Culture supernatant was used for IL-1β ELISA **G**. THP-1 cells were stimulated with PMA as in E and lysed with 0.09% Triton X-100. IL-1β lysates were diluted 1:5 in assay buffer and measured by ELISA. Data represented as mean ± SEM for a minimum of three independent experiments, unless otherwise mentioned. **p<0.01, ***p<0.001 for comparison with respective untreated controls; two-way ANOVA followed by Dunnett’s (A) multiple comparison tests. # p<0.05 Student’s t-test for comparison between colchicine treated cells stimulated with or without 5 μM nigericin.

Prevailing dogma equates speck assembly and inflammasome activation of caspase-1. However, increased speck frequency does not correlate well with levels of released IL-1β. For example, PMA-treated THP-1 macrophages stimulated with 5 μM nigericin yielded approximately 3-fold more speck containing cells than those stimulated with 1 μM despite comparable IL-1β release (data not shown). This lack of correlation between specks and the magnitude of the IL-1β response together with the near complete blockade of IL-1β when colchicine is used with lower doses of nigericin, suggests the possibility that ASC specks might improve, or even maximize, caspase-1 activation by suboptimal stimuli. If correct, the IL-1β response to nigericin is expected to be dose-dependent in the absence of specks. PMA-treated THP-1 macrophages were stimulated with colchicine and stimulated with increasing concentrations of nigericin (1.0 to 7.5 µM). In the presence of colchicine, specks were not observed in cells treated with any concentration of nigericin **(Fig. 1D)** and little IL-1β was produced after stimulation with up to 2.5 μM nigericin (**Fig. 1E**). Interestingly, in the absence of specks (i.e. with colchicine), supernatant IL-1β levels were dependent upon the dose of nigericin for concentrations between 2.5 and 7.5 μM. Similar dose-dependency was observed with LPS-primed THP-1 **(Fig. 1F)**. In the presence of colchicine, higher concentrations of ATP and MSU induced cell death and were not evaluated further. We confirmed that the IL-1β ELISA kit used does not detect uncleaved pro-IL-1β **(Fig. 1G)** released upon cell death. Thus, only IL-1β actively cleaved following stimulation is detected in these assays. While these data demonstrate that the speck is not required for caspase-1 cleavage of IL-1β, they do not rule out the ASC speck as a site of caspase-1 activity. Further, these data also suggest that ASC specks facilitate and maximize caspase-1 cleavage of IL-1β, perhaps through lowering the agonist threshold required for NLRP3 inflammasome activation.

### Caspase-1 activity is distal to the ASC speck organization site

As IL-1β processing can occur in the absence of ASC specks, the site of caspase-1 activation may be distinct from the speck structure when one is present. We previously demonstrated that ASC specks and caspase-1 activity can be visualized simultaneously in cells using imaging flow cytometry (Nagar et al., 2019). Although our focus was on cells with specks and organized caspase-1 activity we observed that active caspase-1 was distributed throughout the cytoplasm. We manually re-evaluated micrographs from these experiments to consider cells with individual specks and active caspase-1 regardless of organization. Inflammasome-reconstituted cells receiving either 20 or 50 ng of procaspase-1 can be divided into four sub-populations based on their distribution of specks and active caspase-1 **(Fig. 2A)**. First, specks with coincident sites of active caspase-1 (Concomitant speck and FLICA (C); 13.5-21.2%). Second, specks and non-coincident (distant) discrete sites of active caspase-1 (Separate speck and FLICA (S); 9.3-9.6%). Third, diffuse caspase-1 activity that overlaps specks (Diffuse FLICA (D); 66.1-66.5%). Four, diffuse caspase-1 activity distributed throughout the cells, but absent at the speck (Non-FLICA specks (N); 24.8-28%) **(Fig. 2B-C)**. Importantly, the amount of transfected pro-caspase-1 did not significantly alter these distribution patterns or the frequency of cells exhibiting each pattern **(Fig. 2B-C)**. Approximately 15-20% of the cells had features of more than one distribution pattern for speck and active caspase-1 and were quantified as overlapping sets **(Fig. 2B-C)**. Populations with ASC specks and organized caspase-1 (S and C) account for 22-30% of the cells **(Fig. 2B-C)**. In population S (2 to 4%), caspase-1 activity was organized and located distally to the ASC speck, suggesting an unexpected spatial disconnect between active caspase-1 and the speck in this subpopulation. Viewing a 3D event in 2D may cause two separate events in distinct planes to appear colocalized. To evaluate colocalization of ASC and active caspase-1 in cells with concomitant staining (population C) using the ImageStream, we used the ‘Bright Detail Similarity’ (BDS) index function and IDEAS-Co-localization Wizard Tool (see Methods). A BDS-index of 3.0 indicates a high degree of colocalization (both objects are in the same focal plane) where an index of 1.0 are not colocalized (not in the same focal plane). Using events from population S as an internal negative control, the BDS-index threshold was set to 1.5. The speck and active-caspase-1 (FLICA) are considered colocalized in cells with scores ≥ 1.5 **(Fig. 2D)**. For nigericin-stimulated cells, the mean BDS-index for the ASC speck and capase-1 activity was approximately 0.25, irrespective of the amount of caspase-1 transfected **(Fig. 2E)** indicating a lack colocalization. Surprisingly, colocalization of the ASC speck and caspase-1 activity (BDS-index ≥ 1.5) occurred in fewer than 2% of cells receiving 50 ng of caspase-1 and none of those receiving 20 ng **(Fig. 2F)**. Thus, even in Population C, where specks and caspase-1 activity appear concomitant, ASC specks and the site of active caspase-1 are spatially separated. Indeed, approximately 90% of cells with specks had diffuse caspase-1 activity (populations D and N) and of these, roughly a third were devoid of caspase-1 activity at the speck itself **(Fig. 2B-C)**. Collectively, these data unexpectedly suggest that in cells with a discrete NLRP3:ASC speck, caspase-1 activity is almost exclusively separate from the speck structure.

**Figure 2:**
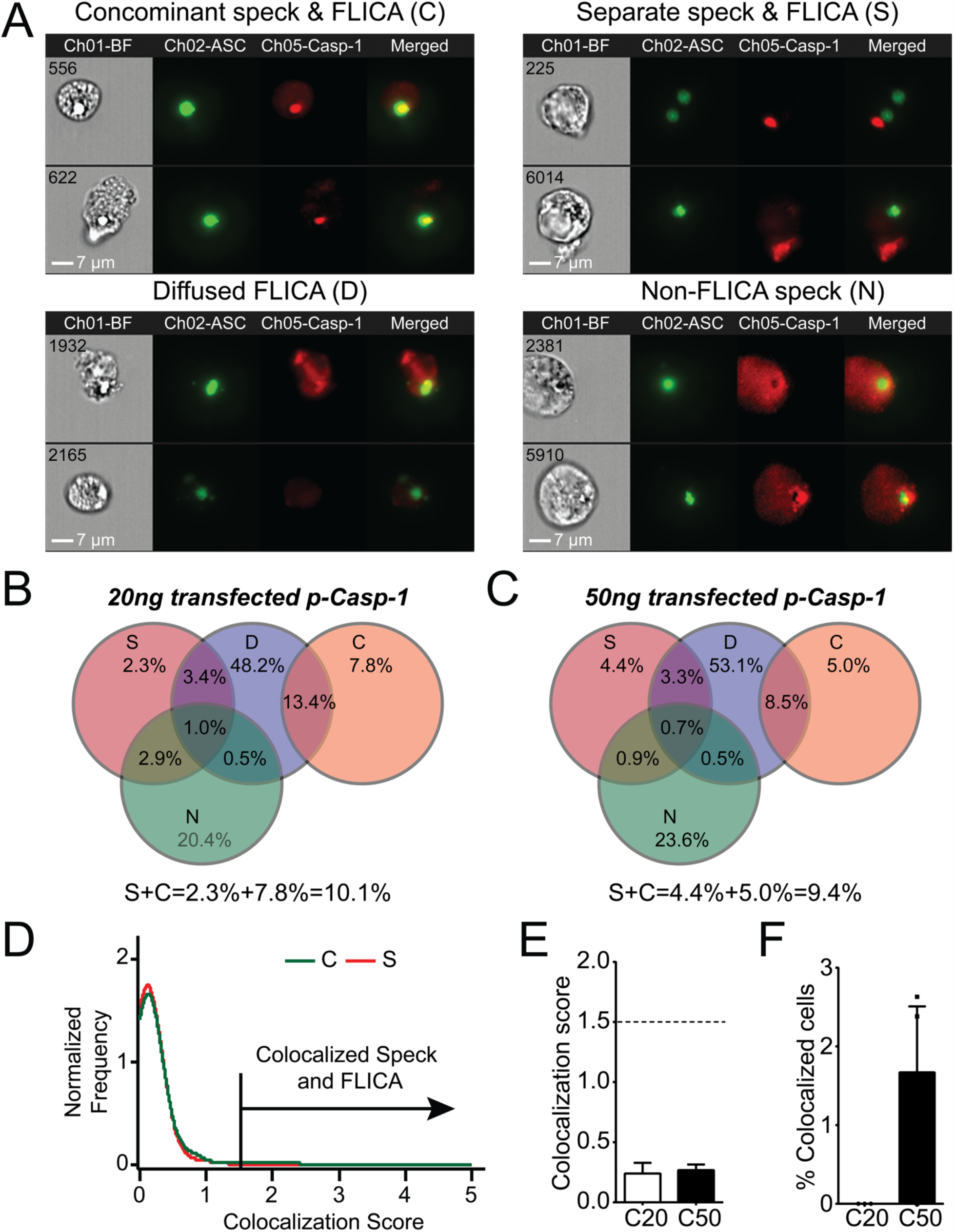
Active caspase-1 does not colocalize with the ASC speck. HEK293T cells were transfected with 100 ng NLRP3, 50 ng GFP-ASC and 20 or 50 ng p-Casp1 and treated with 5 µm nigericin for 2 hrs. ASC speck positive cells were analyzed for distribution of active caspase-1 and ASC speck by ICCE. Ch01-BF (Brightfield), Ch02-GFP-ASC (Green), Ch05-Active caspase-1 (FLICA; red). The cells containing specks and active caspase-1 were manually segregated into four different sub-populations. **A**. Representative images from the sub-population of cells showing (top left) coincident ASC speck and active caspase-1 (C), (top right) distant ASC speck and active caspase-1 (S), (bottom left) diffused active caspase-1 (D) and (bottom right) specks lacking any detectable caspase-1 activity (N). The sub-population of cells where active caspase-1 is diffused around the ASC speck. **B**. Venn diagram showing the frequency of different sub-populations in cells transfected with 20 ng p-Caspase-1 (C20). **C**. Venn diagram showing the frequency of different sub-populations in cells transfected with 50 ng p-Caspase-1 (C50). **D**. Colocalization analysis of ASC speck and active caspase-1 of images shown in A (Subpopulation C), Histogram showing the threshold criteria for colocalization assay. **E**. Mean colocalization score of images shown in A, a threshold score of 1.5 was selected as positive for colocalization. **F**. Frequency of colocalized events over total events noted above the threshold line for C20 (20 ng) and C50 (50 ng) amount of transfected p-Casp-1. Data represented as mean ± SEM for a minimum of three independent experiments. Data represented as Mean for B.

### Speck formation is not equivalent to IL-1β processing

Our analysis suggests the NLRP3:ASC speck and the inflammasome may be distinct entities. Alternatively, if the speck is the site of caspase-1 activity, speck formation should be proportional to IL-1β production. While studies demonstrate that small molecule NLRP3 inhibitors or mutations in inflammasome components block both specks and IL-1β maturation (Coll et al., 2015; Dick et al., 2016; Lu et al., 2014) we found no study reporting proportionality between speck frequency and IL-1β maturation. NLRP3 contains a C8:C108 disulfide bond suggested to be important for NLRP3-ASC interaction (Bae and Park, 2011). Individually, inflammasome activation does not strictly require cysteine 8 or 108, but IL-1β production is increased by C108A and C108S single point mutants and decreased by C8/108 double mutants (Hafner-Bratkovic et al., 2018; Rahman et al., 2020). Since ASC speck formation by these mutants is expected to reflect their inflammasome activity, we evaluated IL-1β release and speck formation in inflammasome-reconstituted HEK293T cells expressing C8 and C108 single and double mutants stimulated with nigericin or infected with *Francisella novicida* (*Fn*) U112. For *Fn* U112 infection, cysteine mutations had little impact on NLRP3-dependent IL-1β maturation or the frequency of ASC speck-containing cells **(Fig. 3A)**. However, with nigericin stimulation IL-1β production increased approximately 1.5X for single-point C108 mutants and decreased by 2X for C8/108 double-mutants relative to wtNLRP3, while speck frequency was diminished by approximately 25% for all C8 and C108 mutants **(Fig. 3B)**. Thus, IL-1β production by C8/108 mutants correlated closely with ASC speck formation (IL-1 β:speck ratio of about 1) in response to Fn U112 infection (**Fig 3C**) but was not correlated when nigericin was used. Indeed, despite all C8/108 mutants having uniformly diminished speck frequency in, single-point C8 mutants decreased nigericin-induced speck formation but had little impact on IL-1β where C108 mutants increased IL-1β production despite diminished speck forming capacity. C8/108 double mutants markedly decreased IL-1β but had little further impact on nigericin-induced ASC speck formation. Thus, speck frequencies do not simply parallel or equate to inflammasome activity (IL-1β production). That IL-1β processing can increase or decrease irrespective of ASC speck formation is consistent with our hypothesis that the site of inflammasome IL-1β processing is distinct from the ASC speck structure.

**Figure 3:**
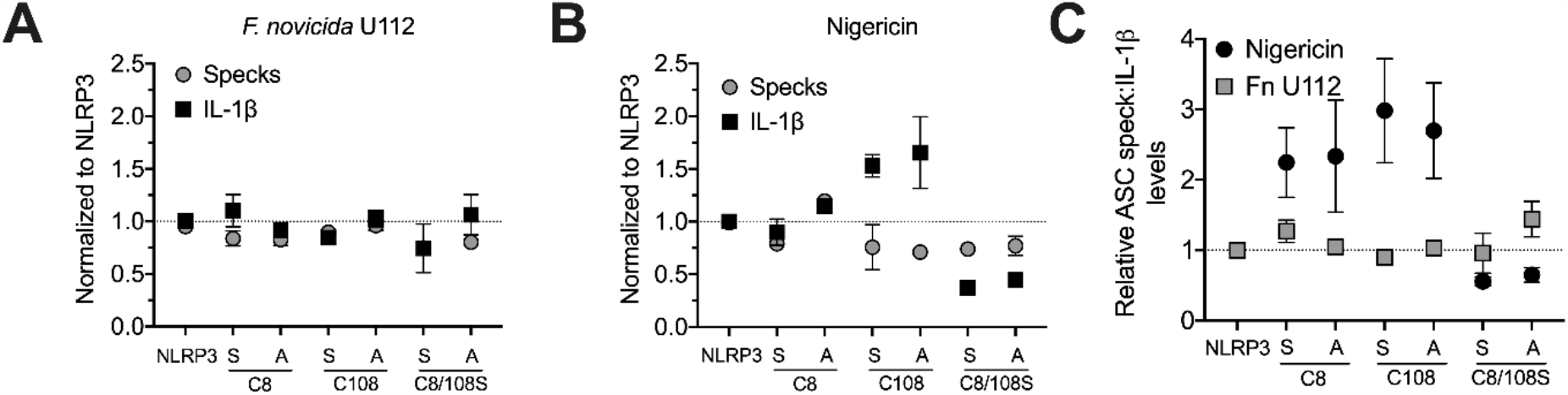
N-terminal cysteine mutations uncouple speck formation and inflammasome function. (For ASC speck analysis) HEK293T were transfected with 50 ng of GFP-ASC with 100 ng NLRP3 or pcDNA3. 18 hrs post-transfection, cells were either stimulated with nigericin agonists or left untreated or 4 hrs after transfection, cells were infected with *Fn*. Cells were fixed at 2 hrs after stimulation (nigericin) or 24 hrs post-*Fn* infection and analyzed for ASC specks by TOFIE. (For IL-1β release) HEK293T were transfected with 8 ng of myc-ASC with 100 ng NLRP3 or pcDNA3, 40 ng p-Casp1 and 200 ng p-IL-1β. 18 hrs post-transfection, cells were either stimulated with indicated agonists or left untreated. **A**. Normalized ASC speck frequency (TOFIE) in comparison to normalized IL-1β release in reconstituted 293T cells following *F*.*n* U112 infection. **B**. Normalized ASC speck frequency (TOFIE) in comparison to normalized IL-1β release in reconstituted 293T cells following nigericin stimulation. **C**. Comparison of the ratio of normalized ASC specks frequency (TOFIE) to normalized IL-1β release following NLRP3 activation by nigericin stimulation and *F*.*n* U112 infection.

Our results with NLRP3 mutants provide compelling evidence that speck formation and inflammasome function are likely disconnected. Although speck formation follows stimulation with inflammasome agonists and precedes IL-1β elaboration, they may not be the site of IL-1β processing. If correct, other indications may be evident during agonist stimulation of wildtype NLRP3. Reconstitution of agonist-responsive NLRP3 inflammasomes is accomplished by transfecting limited quantities of inflammasome components encoding plasmids (Shi et al., 2016). In such experiments, ASC specks are rarely observed beyond background levels despite robust IL-1β production comparable to similarly stimulated macrophages (data not shown). Indeed, to evaluate agonist-dependent induction of specks similar to that seen in macrophages, higher expression of NLRP3 and ASC is required (Nagar et al., 2019; Sester et al., 2015; Shi et al., 2016). Therefore, we compared ASC speck formation capacity (Time of Flight Inflammasome evaluation (TOFIE)) and IL-1β processing (inflammasome reconstitution assay) in HEK293Ts in response to agonists thought to use different activation pathways including pore formation/K^+^ efflux (nigericin), phagosome rupture (*Fn* infection and MSU stimulation), and ROS generation (H_2_O_2_). In cells expressing NLRP3, nigericin treatment induced a 15-fold increase in IL-1β processing and an approximately 3-fold increase in speck formation **(Fig. 4A)**. As expected, no significant speck formation or IL-1β production occurred in unstimulated cells or those lacking NLRP3. Thus, speck formation and IL-1β processing are positively correlated for nigericin stimulation. However, *Fn* infection induced speck formation in the absence of NLRP3, but no IL-1β processing above background was observed without **(Fig. 4B)**. This result is consistent with previous work reporting that specks form in the absence of NLRP3 but lack caspase-1 activity (Compan et al., 2015). In cells expressing NLRP3, *Fn* infection induced a robust IL-1β release without a significant further increase in speck frequency **(Fig. 4B)**. Thus, for *Fn* infection, IL-1β production (inflammasome function) appears disconnected from speck formation. *Fn* induced inflammasome function, but not speck formation, is dependent on NLRP3. Surprisingly, neither H_2_O_2_ nor MSU induced speck formation but significant levels of IL-1β were produced **(Fig. 4C&D)**. In these experiments, approximately 25% of the cells contained a speck, but no IL-1β was produced above controls without stimulation **(Fig. 4C&D)**. For H_2_O_2_ and MSU, speck formation does not equate directly with NLRP3-inflammasome activation. Thus, in inflammasome reconstituted cells, agonists like H_2_O_2_ and MSU may activate the NLRP3 inflammasome without inducing specks.

**Figure 4:**
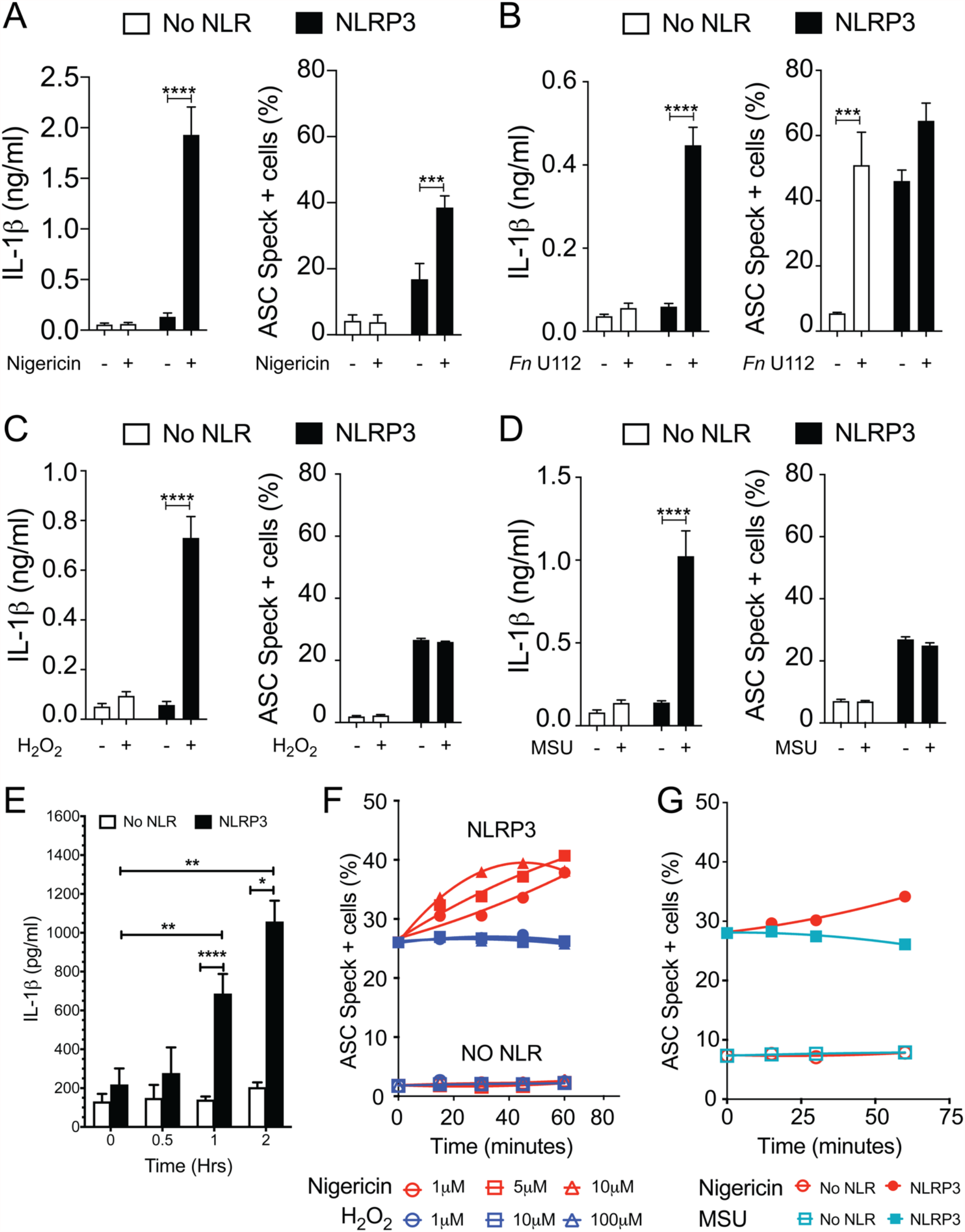
MSU and H_2_O_2_ elicit IL-1β without inducing ASC speck formation. Cells were transfected as described in Fig. 3 and stimulated for NLRP3 activation (as described in Methods: Inflammasome activation) **A**. IL-1β processing and speck formation exhibits positive correlation in nigericin stimulated cells. **B**. Unlike speck formation, IL-1β processing is dependent on NLRP3 in *F. novicida* U112 infected cells. **C**. IL-1β and speck formation shows no correlation in H_2_O_2_ treated cells. H_2_O_2_ treated cells does not induce speck formation. **D**. IL-1β and speck formation shows no correlation in MSU treated cells. MSU treated cells does not induce speck formation. **E**. Time-course analysis of H_2_O_2_ treated IL-1β processing in inflammasome-reconstituted HEK293T cells. **F**. Cells were treated with 5 µm nigericin (red) or 100 µM H_2_O_2_ (blue) for 1 hr and analyzed for ASC speck+ cells by TOFIE. **G**. Cells were treated with either 5 µm nigericin (red) or 150 µg/ml MSU (cyan) for 1 hr and analyzed for ASC speck+ cells by TOFIE. Data represented as mean ± SEM for a minimum of three independent experiments (A-E). Data represented as mean ± SEM for a two independent experiments (F&G). *p<0.05, **p<0.01, ***p<0.001, ****p<0.0001 for comparison with respective untreated, two-way ANOVA followed by Sidak’s multiple comparison tests

As our H_2_O_2_ and MSU results were surprising, we considered that specks induced by these agonists may have been formed but rapidly degraded prior to enumeration. To evaluate this alternate hypothesis, speck formation was tracked for 60 minutes using increasing concentrations of H_2_O_2_ or MSU in cells expressing NLRP3 and ASC. H_2_O_2_ induces IL-1β approximately 60 minutes after stimulation **(Fig. 4E)**. As expected, speck formation increased over time and with escalating nigericin stimulation **(Fig. 4F)**. Interestingly, no concentration of H_2_O_2_ tested induced speck formation above basal levels within 60 minutes and no decrease in basal speck frequency was evident **(Fig. 4F)**. Results with MSU stimulation were essentially identical **(Fig 4G)**. Consistent with the transfected cell data, neither H_2_O_2_ nor MSU induced specks in THP-1 cells **(Fig. S4)**. Thus, rapid speck formation followed by degradation does not account for the absence of specks following H_2_O_2_ or MSU stimulation. Collectively, this data suggests that even when wildtype NLRP3 is considered, speck formation is not equivalent to IL-1β processing and thus the inflammasome may be distinct from the speck.

### Specks are dynamic and negatively correlated with IL-1β release

Our data above strongly suggest that the speck is not necessary for inflammasome activity and that specks may function to instead facilitate inflammasome formation. As active caspase-1 is not localized to the speck, it follows that inflammasome complexes might form outside the speck or alternately be shed from the speck upon caspase-1 activation. Since both the speck and the inflammasome require NLRP3:ASC interaction, envisioning how to demonstrate formation of inflammasome structures outside the speck or shedding of such complexes is problematic. If inflammasomes form exclusively outside the speck, the speck is expected be a static structure. In contrast, if the speck is shedding inflammasomes, its structure should be dynamic (e.g. size may be altered with time and/or activation). To evaluate changes in ASC speck size during NLRP3 activation, we measured speck area in nigericin stimulated cells expressing GFP-ASC, FLAG-NLRP3 and pro-Caspase-1 using our imaging flowcytometry-based inflammasome and caspase-1 activity characterization and evaluation (ICCE) assay (Nagar et al., 2019). Both nigericin treatment or increasing the amount of available pro-caspase-1 significantly decreased speck area **(Fig. 5A)**. The frequency of ASC speck-containing cells also decreased with increased caspase-1 expression in the absence of stimulation **(Fig. 5B)**. These inverse relationships demonstrate that caspase-1 activity negatively regulates both speck area and speck formation and strongly suggest that ASC specks are dynamic in nature.

**Figure 5:**
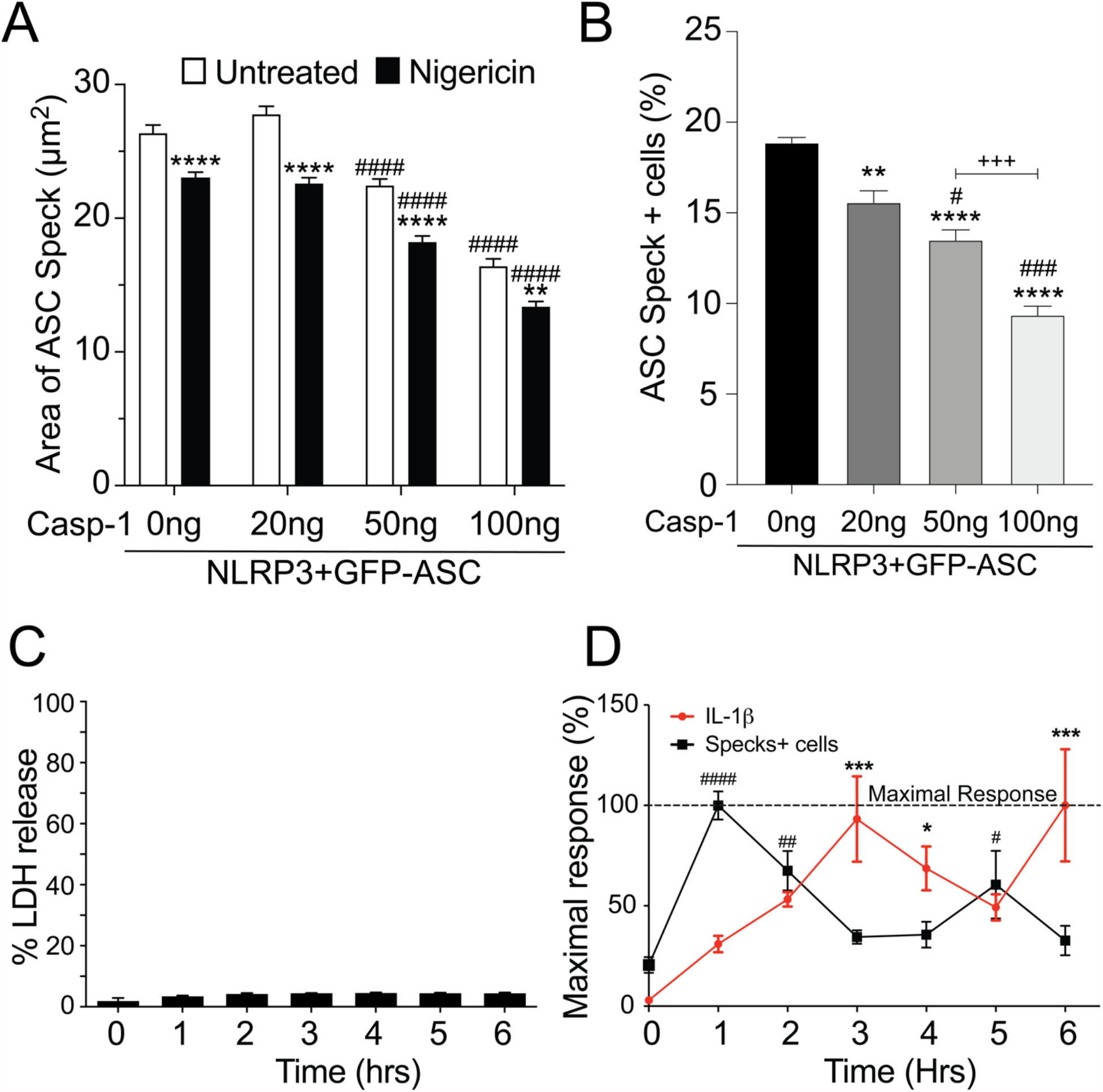
Speck formation and IL-1β release are cyclic and negatively correlated. **(A-B)** HEK293T cells were transfected with 50 ng GFP-ASC, 100 ng NLRP3 and 20 ng or 50 ng or 100ng pCasp-1 and treated with 5 µm nigericin for 2 hrs. Following nigericin treatment, cells were stained with FLICA, fixed and analyzed for the presence of speck and active caspase-1 by ICCE. **(C-D)** PMA differentiated THP-1 cells are either left untreated (0 hrs) or treated with 5 μM nigericin for 6 hrs. Culture supernatant was harvested at stated time-intervals to measure IL-1β by ELISA and release of LDH. Cells were fixed and stained for ASC to detect specks. **A**. Cells were analyzed for the presence of ASC speck and active caspase-1 (positive for FLICA staining) from the double positive gates as analyzed by ICCE. The area of the speck mask was calculated by ICCE. Data represented as mean ± SEM for a minimum of three independent experiments. **p<0.01, ****p<0.0001 for comparison with respective untreated, two-way ANOVA followed by Sidak’s multiple comparison tests. ####p<0.0001 for comparison with respective untreated, two-way ANOVA followed by Dunnett’s multiple comparison tests. **B**. Cells positive for GFP-ASC were analyzed for presence of ASC specks by ICCE. Data represented as mean ± SEM for a minimum of three independent experiments. **p<0.01, ****p<0.0001 for comparison with 0 ng transfected casp-1, #p<0.05, ###p<0.001 for comparison with 50 ng transfected casp-1, +++ p<0.001 for comparison between 50 ng and 100 ng transfected casp-1, one-way ANOVA followed by Tukey’s multiple comparison tests. **C**. Percentage LDH release compared to 0.09% Triton-X100-lysed cells. **D**. Line graph showing the frequency of ASC speck+ cells (red) and release IL-1β (black). Data is normalized to maximal response of speck frequency and released IL-1β over the time course of the experiment. Data represented as mean ± SEM for a minimum of three independent experiments. *p<0.05, ***p<0.001, for comparison of IL-1β release with untreated control (0 hrs), #p<0.05, ##p<0.01, ####p<0.0001 for comparison of speck frequency with untreated control (0 hrs), one-way ANOVA followed by Sidak’s multiple comparison tests.

Since caspase-1 activity was negatively correlated with speck frequency in transfected cells, we also examined this relationship in macrophages. Nigericin was selected because neither H_2_O_2_ or MSU induced specks in THP-1 cells **(Fig. S3)** and nigericin-induced speck formation appears to be directly correlated with IL-1β processing **(Fig. 4A)**. No significant LDH release was detected over a 6 hr time-course **(Fig. 5C)**. PMA-primed THP-1 cells were activated with nigericin and speck formation and IL-1β release were followed for 6 hrs; beyond 6 hrs cells begin to detach from the plate complicating assessment of speck formation. Specks were induced by nigericin, but their frequency was cyclical over time **(Fig. 5D)** and reached maximum around 1 hr. Speck frequency declined to near basal levels over the next 2 hrs, increased to about 50% of the maximum by 5 hrs post-stimulation, and then again decreased. IL-1β release was also cyclical. IL-1β levels increased over time but did not reach a maximum until around 3 hrs. By 5 hrs, IL-1β levels had declined to about 50% of the maximum before increasing to near maximum levels around 6 hours. Importantly, ASC specks were not detected outside the cells (data not shown). While both speck formation (frequency) and IL-1β were cyclic in nature, they were out of phase, with IL-1β increasing after peaks of speck formation. Although specks precede IL-1β production, consistent with specks facilitating production of IL-1β, the temporal lag between speck frequency and IL-1β levels (2 hrs for the first cycle) reveals that peak caspase-1 activity likely follows well after peak speck formation (see **Discussion**). Under in vitro conditions, the Vmax of caspase-1 in an NLR complex is around 2 nM/minute (Faustin et al., 2007). At this rate, the quantity of IL-1β present at 3 hrs (5 ng/ml; ∼0.3 nM) should have been converted in less than one minute, i.e. with no temporal lag. Thus, it is likely that IL-1β processing does not occur simultaneously with formation of the speck, but instead develops gradually over time following speck formation.

### Small inflammasome complexes are shed from the speck

The physical and temporal separation of caspase-1 activity from specks suggests that inactive NLRP3:ASC:caspase-1 complexes might be shed from the speck with activation of caspase-1 and processing of IL-1β occurring sometime later. If correct, cells with multiple sites of caspase-1 activity (with or without some diffuse activity) are expected to be abundant. Therefore, we evaluated the frequency of such cells in our HEK293T data from Figure 2. Nigericin-stimulated primary human macrophages and immortalized BMDMs were also examined. Multiple FLICA stained sites indicating caspase-1 activity were observed in approximately 30% of speck-containing reconstituted HEK293T cells **(Fig. 6A)**, 55-60% of speck-containing primary human macrophages stimulated with nigericin or ATP **(Fig. 6B)** and 40% of nigericin-stimulated immortalized BMDMs **(Fig. 6C)**. While not direct evidence, the substantial number of cells with the expected phenotype is consistent with the shedding of inflammasome complexes from the speck. Speck-independent activation of small inflammasome complexes could also explain the observed pattern of caspase-1 activity. Indeed, small “death complexes” have been implicated in pyroptosis (caspase-1-dependent cell death) and IL-1β processing after Aim2 and NLRC4 inflammasome activation in macrophages (Broz et al., 2010; Dick et al., 2016).

**Figure 6:**
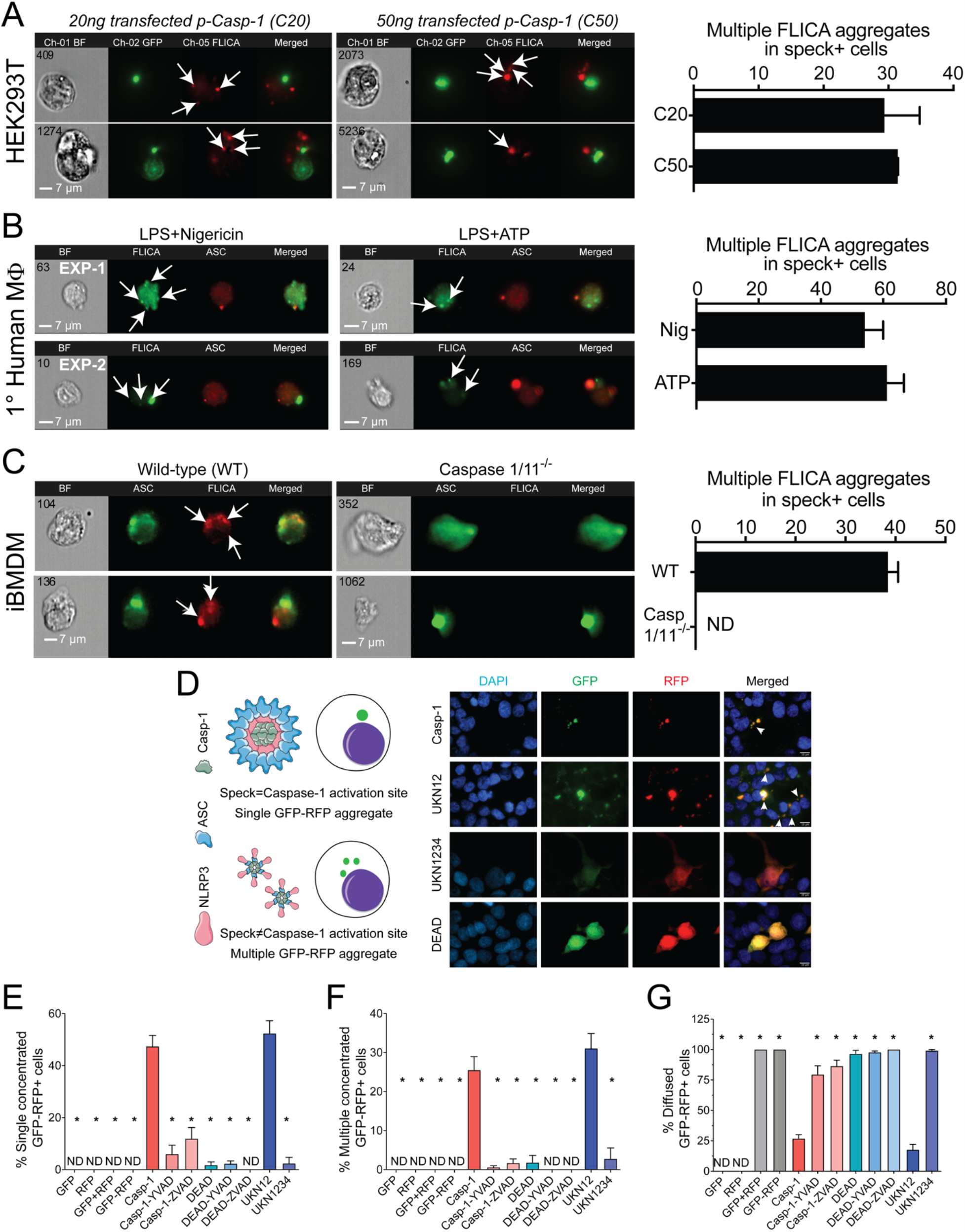
NLRP3-dependent caspase-1 activity occurs at multiple cellular sites. LPS primed primary human monocytes and immortalized BMDMs cells were either left untreated or stimulated with 5 mM ATP or 10 µm nigericin for 30 mins, stained with FLICA and were fixed and analyzed by ICCE. HEK293T cells were transfected with 50 ng GFP-ASC with or without 100 ng NLRP3 and stimulated with 5 μM nigericin. Following nigericin stimulation cells were stained with FLICA, fixed and analyzed by ICCE. **A**. (Left) The ASC (green) speck+ cells containing active caspase-(red) showed multiple FLICA (active caspase-1) aggregates in both cells transfected with 20 ng and 50 ng of p-Caspase-1. (Right) The frequency of cells containing speck and multiple FLICA aggregates. Data represented as mean ± SEM. **B**. (Left) Primary human macrophages containing ASC speck (red) showing FLICA staining for active caspase-1 (green) as diffused and/or aggregated (multiple and distant from the speck). Representative images from 2 independent experiments are shown. (Right) The frequency of speck+ cells containing multiple FLICA aggregates. Data represented as mean ± SEM. **C**. (Left) Immortalized BMDMs containing ASC speck (green) showing FLICA staining for active caspase-1 (red) as diffused and/or distant aggregates from ASC speck. Caspase-1/11^-/-^ BMDMs shows no FLICA staining showing specificity of staining. (Right) The frequency of cells containing speck and multiple FLICA aggregates. Data represented as mean ± SEM. **(D-G)** HEK293T cells were transfected with 100 ng myc-ASC, 100 ng NLRP3, 400 ng p-Casp1 (WT or mutant) and 400 ng GFP-YVAD-RFP. 18 hrs post-transfection, cells were fixed analyzed for GFP-RFP aggregates by microscopy. **D**. Schematic diagram showing expected outcomes with bi-fluorescent reporter if single/multiple sites of active caspase-1 are present. Representative micrographs showing distribution of active caspase-1 sites in a cell; DAPI (blue), GFP (Green), RFP (Red) and Merged. White arrows show cells with multiple GFP-RFP aggregates **E**. Frequency of cells containing single concentrated GFP-RFP aggregates. **F**. Frequency of cells containing multiple concentrated GFP-RFP aggregates. **G**. Frequency of cells containing diffused GFP-RFP. Data represented as Mean±SEM for each field of view for at least three independent experiments. A minimum of 1000 cells were analyzed for each condition. *p<0.0001, for comparison with casp-1 transfected sample, one-way ANOVA followed by Dunnett’s multiple comparison tests.

To better distinguish between these possibilities, we attempted to address whether caspase-1 is activated in the vicinity of the speck or at some distance using various caspase-1 mutants **(Fig. S2A)**. Cleaved caspase-1 p35/p10 is the active form and remains complexed with NLRP3:ASC until further cleavage generates inactive p10/p20 tetramers which leave the complex (Boucher et al., 2018). Mutation of caspase-1 aspartic acids (D) 103 and 119 to asparagine (N) in caspase-1 UKN12 prevents inactivating cleavage events and the dissociation of active caspase-1 (p35/p10) from the NLRP3:ASC complex. UKN1234 has mutations (D→N) in all four aspartic acid cleavage sites preventing its activation. Mutation of caspase-1 cysteine 285 to alanine alters the active site reducing activity (DEAD). Although FLICA staining appears to be caspase-1 specific **(Fig. 6C)**, others have suggested that FLICA reagents are promiscuous (Darzynkiewicz and Pozarowski, 2007). Thus, for these experiments, the FRET-reporter GFP-YVAD-RFP was used to detect active caspase-1 (See Methods; **Fig. S1**). Wildtype and mutant caspase-1 activity measured using the GFP-YVAD-RFP reporter was comparable to that reflected by secreted IL-1β levels, suggesting similar kinetics and specificity for both methods **(Fig. S2B-C)**. Localization of GFP-RFP signal to a single perinuclear complex is expected if caspase-1 activation is occurring at the ASC speck. In contrast, multiple GFP-RFP aggregates would indicate that caspase-1 is activated outside the speck **(Fig. 6D)**. Active inflammasomes were reconstituted in HEK293T cells expressing wildtype or mutant caspase-1 in the presence of the GFP-YVAD-RFP reporter and evaluated for their GFP-RFP staining pattern by wide-field microscopy (single, multiple, or diffuse) **(Fig. 6D)**. No aggregates were observed in cells transfected with GFP-alone, RFP-alone, GFP with RFP, or the GFP-YVAD-RFP reporter **(Fig. 6E&F)**, indicating that these constructs do not spontaneously aggregate. With wildtype caspase-1, multiple GFP-YVAD-RFP complexes were present in 25% of the cells, while 50% had a single complex, and 25% had diffuse staining **(Fig. 6D-G)**. Using the activatable, non-released caspase-1 mutant UKN12 did not alter these frequencies. Both types of complexes were absent when caspase-1 was inhibited with a caspase-1 specific inhibitor (YVAD) or a pan-caspase-1 inhibitor (ZVAD) and when the enzymatically inactive caspase-1 DEAD or uncleavable caspase-1 mutant UKN1234 were used. Some caspase-1 transfected cells had single speck-like GFP-RFP aggregates **(Fig. S4A)**. The absence of active caspase-1 co-localization with specks **(Fig. 2E&F)** suggests this activity is near, but not coincident with, the ASC speck. In caspase-1-UKN12 transfected cells, a zone of no GFP-RFP staining was observed at the likely position of the speck **(Fig. S4B)** similar to the Non-FLICA Speck (N) cell population **(Fig. 2A)**. That UKN12 appears inactive at the likely position of the speck suggests that caspase-1 is activated after leaving the speck structure. Taken together, our data suggests that caspase-1 is most likely activated upon leaving the speck within small inflammasome complexes.

## DISCUSSION

### Caspase-1 is activated outside the speck

Our data provides the first clear demonstration that the speck is not the site of NLRP3-mediated caspase-1 activity. Instead, in both transfected cells and primary macrophages NLRP3 activated caspase-1 is almost exclusively present in multiple cytoplasmic aggregates. This observation seemingly contradicts two studies suggesting that caspase-1 is activated at the NLRP3-ASC speck complex. In one, active caspase-1 was co-isolated with chemically crosslinked ASC (Boucher et al., 2018). However, whether the caspase-1 activity observed in such studies was isolated from the speck itself or some other smaller complexes is uncertain as ASC crosslinking does not distinguish between these complexes. In the other study, active caspase-1 was colocalized to the center of the speck structure formed during NLRC4:NLRP3 inflammasome activation (Man et al., 2014). Although we observed some cells with a similar coincident pattern, further analysis indicates vertical separation of the NLRP3:ASC speck and active caspase-1. In contrast to these studies, we observed that NLRP3 directed caspase-1 activity localized within multiple small cytoplasmic complexes. Consistent with our results, Akira et al also observed multiple NLRP3-ASC complexes in IL-1β producing cells stimulated with monosodium urate crystals using proximity ligation assay (Misawa et al., 2013), although the absence of a speck in these assays was not mentioned or discussed. Further, we also observed this localization pattern with the caspase-1 mutant UKN12. As UKN12 cannot be released from the inflammasome complex upon activation (Boucher et al., 2018), this data confirms the presence of multiple cytoplasmic inflammasome complexes distinct from the speck.

### Speck formation is not essential or sufficient for inflammasome response

Formation of the singular ASC speck is very rapid, occurring within 3 mins of NLRP3 activation (Franklin et al., 2014; Stutz et al., 2013). In addition, inhibiting speck assembly using small-molecule drugs (e.g. MCC950) or Pyrin-related peptides (e.g. POP2) also impairs NLRP3 inflammasome activity (Atianand and Harton, 2011; Coll et al., 2015). Such observations have been interpreted to indicate that the speck is equivalent to the inflammasome or, at least, an initial step in inflammasome activation. However, despite this strong correlative evidence, no direct experimentation has confirmed that the speck is actually the inflammasome.

We previously noted that nigericin activates caspase-1 without discernable ASC speck formation in NLRP3 inflammasome-reconstituted HEK293T cells (Nagar et al., 2019), leading us to question the requirement of specks for inflammasome function. In the present study, the sterile agonists MSU and H_2_O_2_ also induce IL-1β processing without speck formation in HEK293T cells. Compan et al observed that stimulation with 200mg/ml of MSU for 16 hrs induced speck formation in BMDMs (Compan et al., 2015). In our hands THP-1 cells only required 2 hrs of stimulation with 150μg/ml MSU to produce IL-1β and speck formation was absent **(Fig. S3)**. Further, the MSU used by Compan et al. was manufactured by a different company. Thus, these differences may account for the contrasting observations between the studies. Nevertheless, if MSU-induced specks form at 16 hrs in THP-1, this appears to be well after the initiation of IL-1β processing. Of note, and as mentioned above, Akira et al observed multiple NLRP3-ASC complexes, as opposed to single specks, upon stimulation with MSU (Misawa et al., 2013). H_2_O_2_ induces inflammasome processing of IL-1β in THP-1 and other cells (Zhou et al., 2010) but no studies report that H_2_O_2_ induced speck formation. In THP-1 cells H_2_O_2_ did not stimulate speck formation **(Fig. S4)**. Indeed, although inducing inflammasome activity, H_2_O_2_ may inhibit speck formation as suggested during *Streptococcus pneumoniae* infection (Erttmann and Gekara, 2019).

Moreover, speck assembly does not necessitate activation of caspase-1. We observed that *Fn* U112 elicits ASC speck in cells lacking NLRP3, but IL-1β processing is not detected. Others report that osmotic changes induce speck formation in the absence of NLRP3, but these cells also do not process IL-1β (Compan et al., 2015). Moreover, chimeric NLRP3 constructs containing the pyrin domain of either NLRP1 or 2 form specks but not an active inflammasome **(Fig. S5)** (Rahman et al., 2020). Of note, reconstitution of speck formation and inflammasome function requires different amounts of transfected ASC. When agonist-dependent inflammasome activation is reconstituted in HEK293T cells, specks are not evident; higher expression of ASC is required to observe agonist-dependent or independent speck formation (Nagar et al., 2019), again supporting the conclusion that ASC speck formation and inflammasome function are distinct. Although surprising given the seemingly exclusive linkage between ASC specks and active inflammasomes, our data demonstrates that formation of an ASC speck is not required for NLRP3-dependent caspase-1 activation, nor is the ASC speck sufficient for caspase-1 activation.

### The speck lowers the agonist threshold of for NLRP3 inflammasome activation

If neither required for NLRP3 inflammasome activation nor sufficient, what is the function of the speck? Given the prion-like polymerization of ASC and caspase-1 along with the defined perinuclear localization of singular specks in cells, the ASC speck has been suggested to be a <u>s</u>upra<u>m</u>olecular organizing center (SMOC) (Kagan, 2012; Kagan et al., 2014; Wu, 2013). SMOCs are cell location-specific higher-order signaling complexes that concentrate signaling components to facilitate binary, all-or-none, responses through a cooperative assembly mechanism (Kagan, 2012; Kagan et al., 2014; Wu, 2013). An ASC SMOC is expected to recruit most if not all of the available ASC and caspase-1 to the structure, maximize proximity-induced activation of caspase-1, and reduce the threshold of activation. Thus, activation of caspase-1 in this fashion should also occur in an essentially binary, “on-off” fashion. Why caspase-1 is not obligately activated upon speck formation is puzzling, but our data offers some insight.

NLRP1 or NLRP2 Pyrin domain chimeras of NLRP3 form specks **(Fig. S5A)** and *Fn* U112 elicits ASC specks in cells lacking NLRP3. Yet, IL-1β processing is either absent or significantly reduced versus controls in both cases **(Fig. S5B)**. The simplest interpretation is that a specific NLR is required to either recruit or activate recruited caspase-1. However, this conflicts with current paradigms. Oligomerization of ASC recruits caspase-1 (Lu et al., 2014) and once recruited, caspase-1 is expected to undergo proximity-induced activation (Salvesen and Dixit, 1999). Paradoxically, our data indicates that NLRP3-dependent caspase-1 activity is found in small cytosolic complexes distinct from the speck. Further, activity of the caspase-1 UKN12 mutant (activatable, non-dissociating) was also not localized to the speck. As UKN12 cannot dissociate from NLRP3:ASC and UKN12, complexes with active UKN12 must have been activated either independently or after release from the speck (Boucher et al., 2018). A potential resolution to this dilemma is that while small, non-speck NLRP3 inflammasome complexes can form without the speck itself, the speck facilitates formation and release of small inflammasome complexes, thereby amplifying or lowering the threshold for inflammasome activation. Consistent with this idea, Dick et al. suggest that the NLRC4 induced ASC speck serves as a signal amplification mechanism through formation of multiple caspase-1 activation sites (Dick et al., 2016). While exploration of this hypothesis is needed, the seminal studies of extracellular inflammasomes/specks do not appear to be contradictory (Baroja-Mazo et al., 2014; Franklin et al., 2014).

Such a mechanism might also help explain the threshold effect of the speck for nigericin-induced NLRP3 inflammasome activation. When speck formation is inhibited with colchicine, low dose nigericin fails to activate the NLRP3 inflammasome, but higher doses of nigericin still elicit IL-1β (Gao et al., 2016; Van Gorp et al., 2016). In our hands, higher doses of nigericin elicited IL-1β maturation without speck assembly. Notably, when speck formation was inhibited by colchicine, nigericin induction of IL-1β was largely dose dependent. In contrast, without colchicine, IL-1β was elaborated at or near maximal levels irrespective of the nigericin concentration used. Thus, while not essential for NLRP3 inflammasome activation and unlikely the site of caspase-1 activity, the speck reduces the threshold required for agonist induction of full caspase-1 activity. Our findings are consistent with the NLRP3:ASC speck having SMOC-like behavior lowering the threshold of inflammasome activation. However, the physical separation of caspase-1 activity from the ASC speck suggests that the NLRP3:ASC SMOC facilitates the formation and release of small, cytosolically active inflammasomes. Although consistent with our data, this hypothetical mechanism requires further investigation.

### The speck is a dynamic structure regulated by caspase-1

Whether the ASC speck is a SMOC that facilitates the formation and release of small inflammasome complexes is unclear and demonstrating this presents numerous challenges. ASC specks are larger in cells with enzymatically inactive caspase-1 (Caspase-1 DEAD) compared to cells with wildtype caspase-1 (Stein et al., 2016). Moreover, ASC specks formed by ASC alone are larger than NLRP3:ASC specks and the size of NLRP3:ASC specks is further reduced after nigericin stimulation (Nagar et al., 2019). In this study, NLRP3:ASC speck size was inversely proportional to caspase-1 activity and nigericin stimulation further reduced the size of these specks. Thus, speck size is dynamic and is regulated by caspase-1 activity likely resulting from triggering of NLRP3. This change in size suggests that NLRP3 activation of caspase-1 may allow shedding of component proteins from the speck and could account for the presence of multiple cytosolic sites of inflammasome caspase-1 activity. While a change in protein conformation and/or spatial organization could account for the reduction in speck size, such changes do not help explain the multiplicity of active capsase-1 sites.

Interestingly, specks and inflammasome function are not only spatially separated, but temporally distinct as well. We found that speck frequency and IL-1β processing were negatively correlated over time. Further, the relationship between specks and IL-1β processing was also cyclical with declines from peak speck frequency preceding IL-1β processing, suggesting an inverse relationship between the speck and inflammasome function. Further, in inflammasome-reconstituted HEK293T cells, abundant IL-1β processing occurred in the relative absence of ASC specks, but prior to their peak (∼7%) at 14 hours **(Fig. S5C-F)**. Curiously, osmotic-stress induced ASC specks in bone marrow-derived macrophages also suggests a similar inverse relationship between the speck and inflammasome activity (Compan et al., 2015). The temporal disconnect between speck formation and inflammasome function provides additional evidence that speck function is dynamic. Moreover, ASC speck formation might not be irreversible as believed.

In summary, we conclude that ASC specks and NLRP3 inflammasomes are distinct entities and that not all NLRP3 inflammasome agonists trigger speck formation. While small active NLRP3 inflammasome complexes are dispersed throughout the cell following activation and caspase-1 activity is not present at the NLRP3:ASC speck, the speck appears to enhance this process by reducing the threshold of stimulus needed for maximal caspase-1 activation. Our data suggests that when specks are formed, NLRP3 inflammasome IL-1β processing and speck size are related in a dynamic and cyclical process regulated by caspase-1. We propose a refined hypothetical model for the relationship between the NLRP3 speck and inflammasome where formation of an NLRP3:ASC:Caspase-1 speck is energetically favored and releases small active inflammasome complexes but is not required to generate them **(Fig. 7)**.

**Figure 7:**
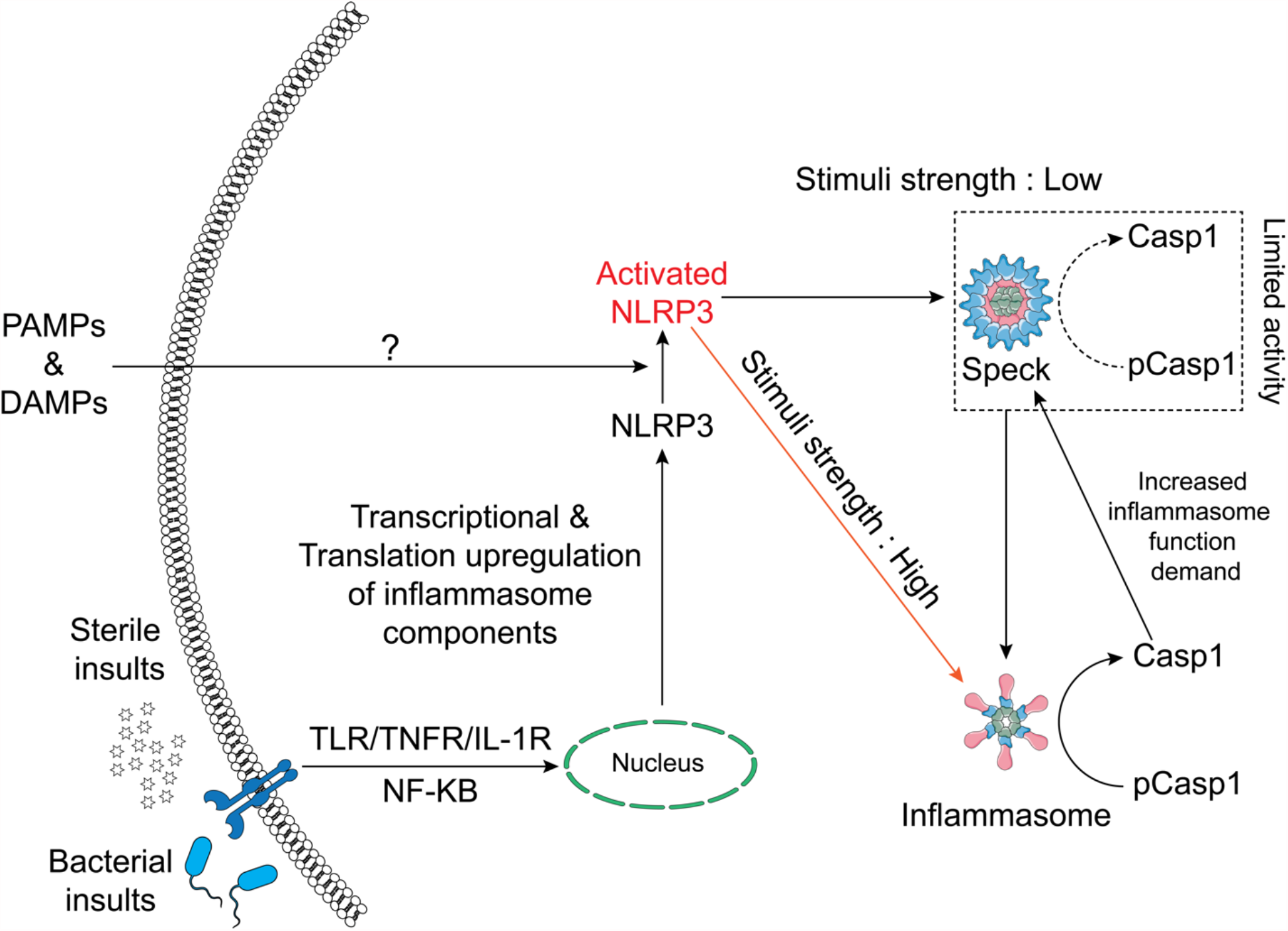
Schematic of the relationship between the NLRP3 inflammasome and speck. NLRP3-ASC specks form by rapid relocation of these proteins into a perinuclear toroidal structure. Specks allow inflammasome complexes to be activated at a lower stimulus threshold and thus facilitate an optimal inflammasome response to weaker signals. However, inflammasome complexes also form independently of the speck and stronger signals may not require or elicit speck formation. The speck likely sheds small inflammasome complexes where caspase-1 becomes active. As caspase-1 activity regulates the size of the speck, active caspase-1 may facilitate the release of small inflammasome complexes, acting as an intracellular feedforward loop maximizing IL-1β processing.

## SUPPLEMENTARY FIGURES

**Figure S1:**
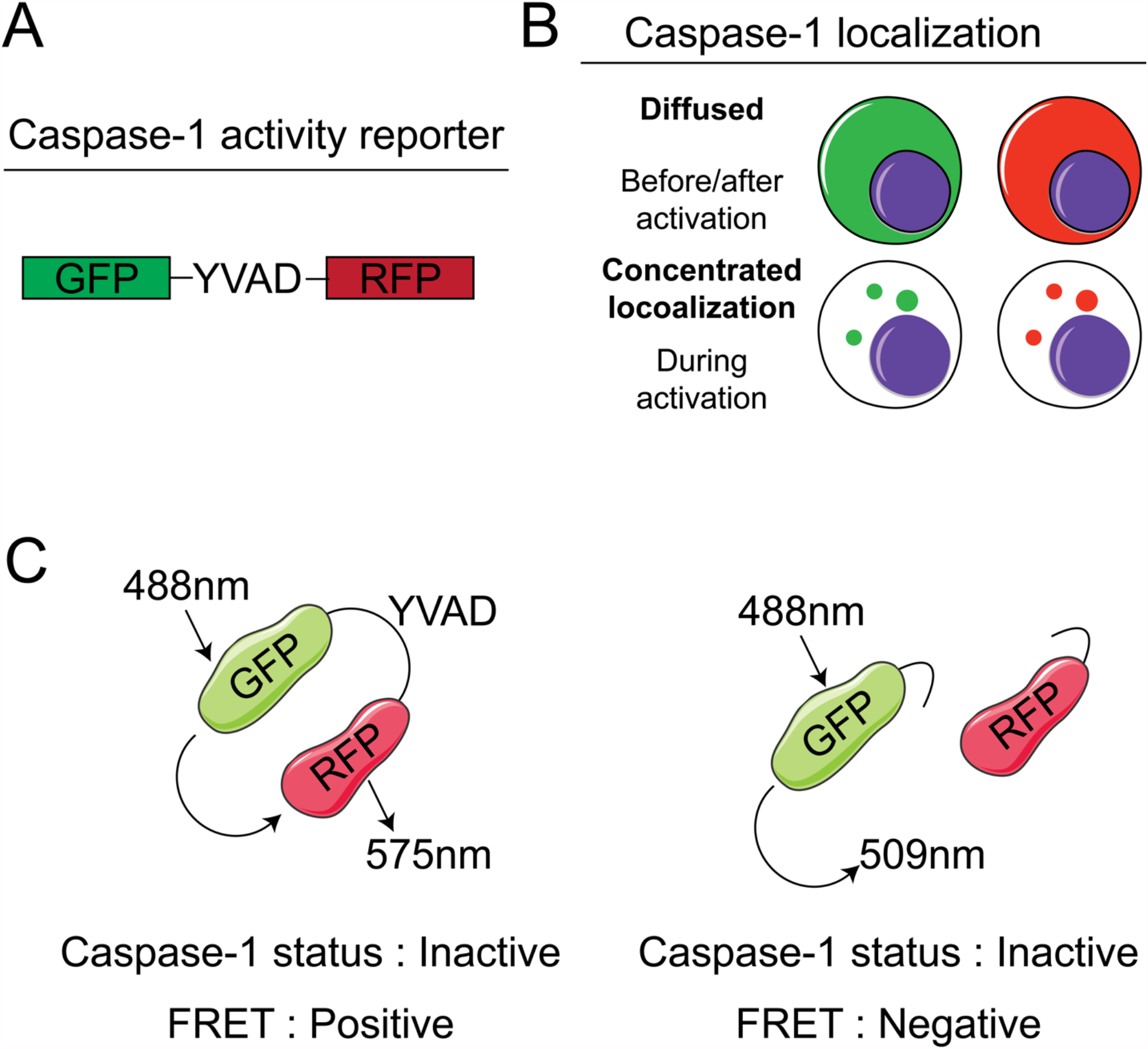
Schematic representation of FRET based bi-fluorescent caspase-1 activity reporter. **A**. Schematic representation of bi-fluorescent reporter for caspase-1 activity. GFP and RFP are linked by the caspase-1-specific recognition amino acid sequence “YVAD”, which is also present in pro-IL-1β. **B**. The expected distribution patterns of caspase-1 reporter before, during, and after caspase-1 activation. **C**. Schematic showing the cleavage states of the bi-fluorescent reporter allowing quantitation of cleavage by caspase-1.

**Figure S2:**
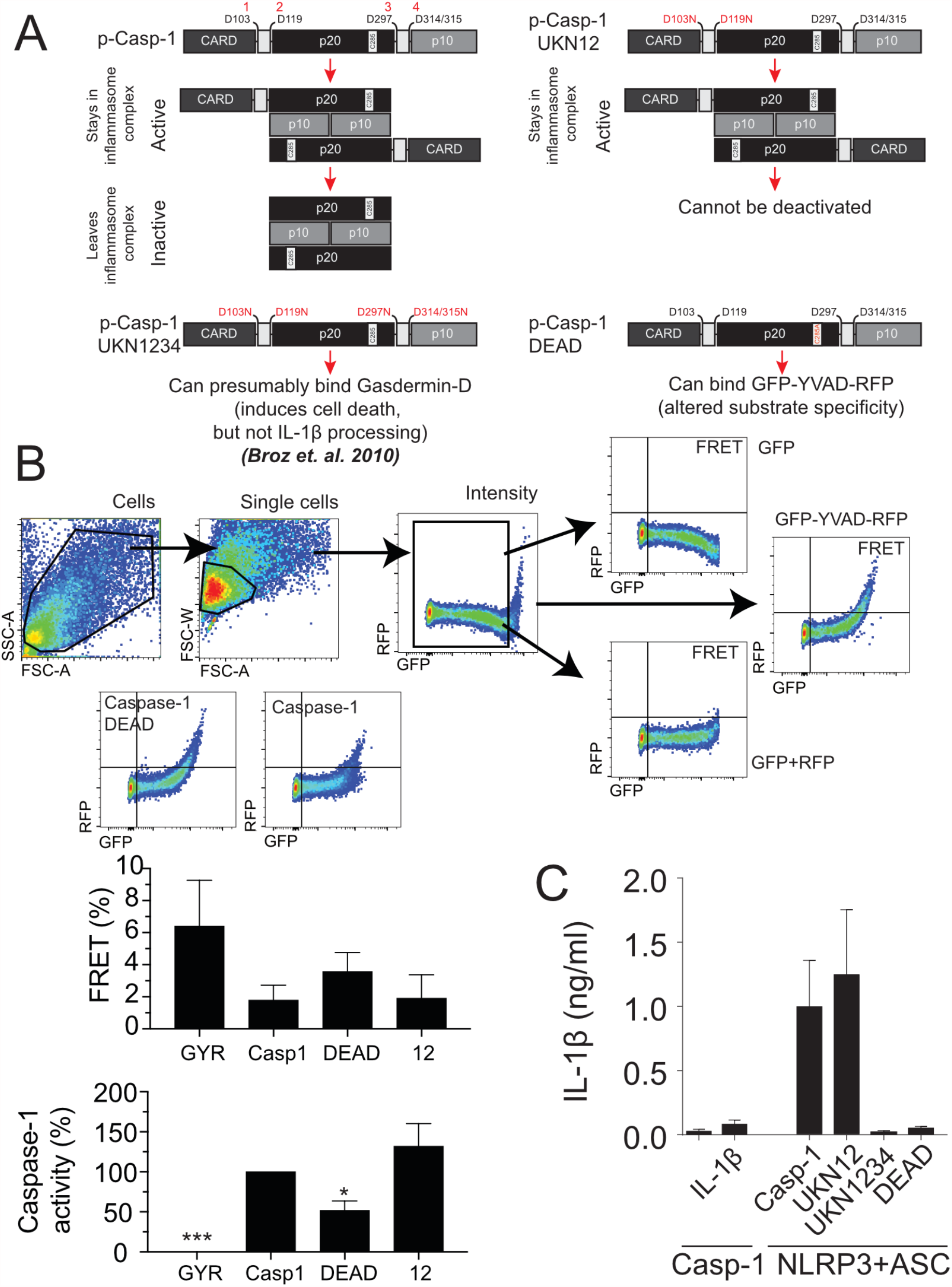
Schematic representation of Caspase-1 mutants, their physiological phenotype and a FRET based bi-fluorescent caspase-1 activity reporter. HEK293T cells were transfected with 100 ng Flag-NLRP3, 100 ng myc-ASC,400 ng pro-IL-1β (B) or GFP-YVAD (D), 400 ng pro-caspase-1 (WT or mutants) or empty vector (EV). **A**. Schematic representation of caspase-1 mutants and their expected phenotypes. The position of five aspartic acid residues are shown (103, 119, 297, 314 and 315). Aspartic acids 103 and 119 are the cleavage sites between the CARD domain and p20 subunit of caspase-1. Aspartic acid residues 297 and 314/315 are the cleavage sites between the p20 and p10 subunit of caspase-1. The caspase-1 UKN12 mutant (D103N/D119N) can be activated but cannot be released or deactivated. All the aspartic acid cleavage sites have in the caspase-1 UKN1234 mutant have D→N mutations, making this mutant non-activatable. The caspase-1 DEAD mutant has the caspase deactivating C285A active site mutation. **B**. The gating strategy to determine caspase-1 activation by flow cytometry. (From left to right: Top) The SSC-A vs FSC-A plot allows to eliminate debris and gate for cells. The FSC-W vs FSC-A plot allows to gate for single cells. The singlets were then plotted for optimum GFP intensity (PE-A vs FITC-A). The optimum GFP intensity cells were then plotted for FRET and a quadrant gate was made to determine FRET positive cells. Representative plots showing the absence of events in FRET channel for cells transfected with GFP-alone or GFP and RFP expressed on different plasmids. Events in FRET channel can be recorded only in cells transfected with GFP-YVAD-RFP. (Bottom) The quadrant plot showing FRET intensity of GFP-YVAD-RFP (bi-fluorescent reporter) in FRET channel of samples transfected with caspase-1 or caspase-1 DEAD. Bar graphs showing % FRET positive cells and % caspase-1 activity (Calculations in Methods and Materials) **C**. 24 hrs post-transfection, culture supernatant was collected and IL-1β was measured by ELISA

**Figure S3:**
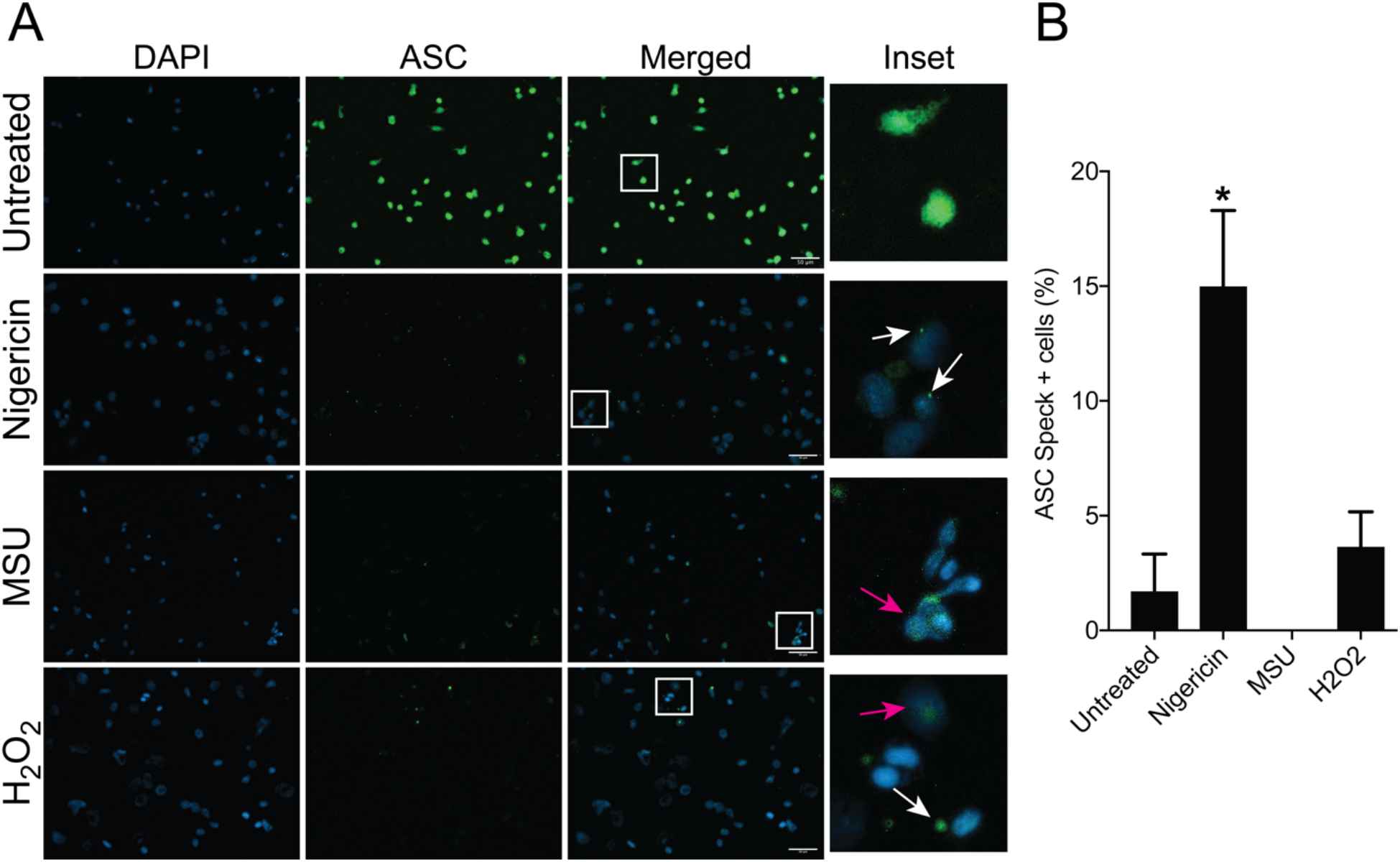
NLRP3 inflammasome activation does not necessitate ASC speck formation. **A**. THP-1 cells were primed with LPS for 4 hrs and stimulated with the stated agonist (See **Methods**) Cells were fixed and permeabilized. Cells were stained for ASC and nuclei were stained with DAPI. Cells were analyzed for presence of specks. Scale bar = 50 µm. White arrows denote specks, yellow arrowhead indicates extracellular specks and red arrow indicates examples of non-speck cells. **B**. Percentage of cells with intracellular specks and extracellular specks from A. Minimum of 100 cells from 3 field of views were analyzed. Mean percentages ± SEM are shown; **p<0.01 & *p<0.5, for comparison with untreated sample, one-way ANOVA followed by Dunnett’s multiple comparison tests.

**Figure S4:**
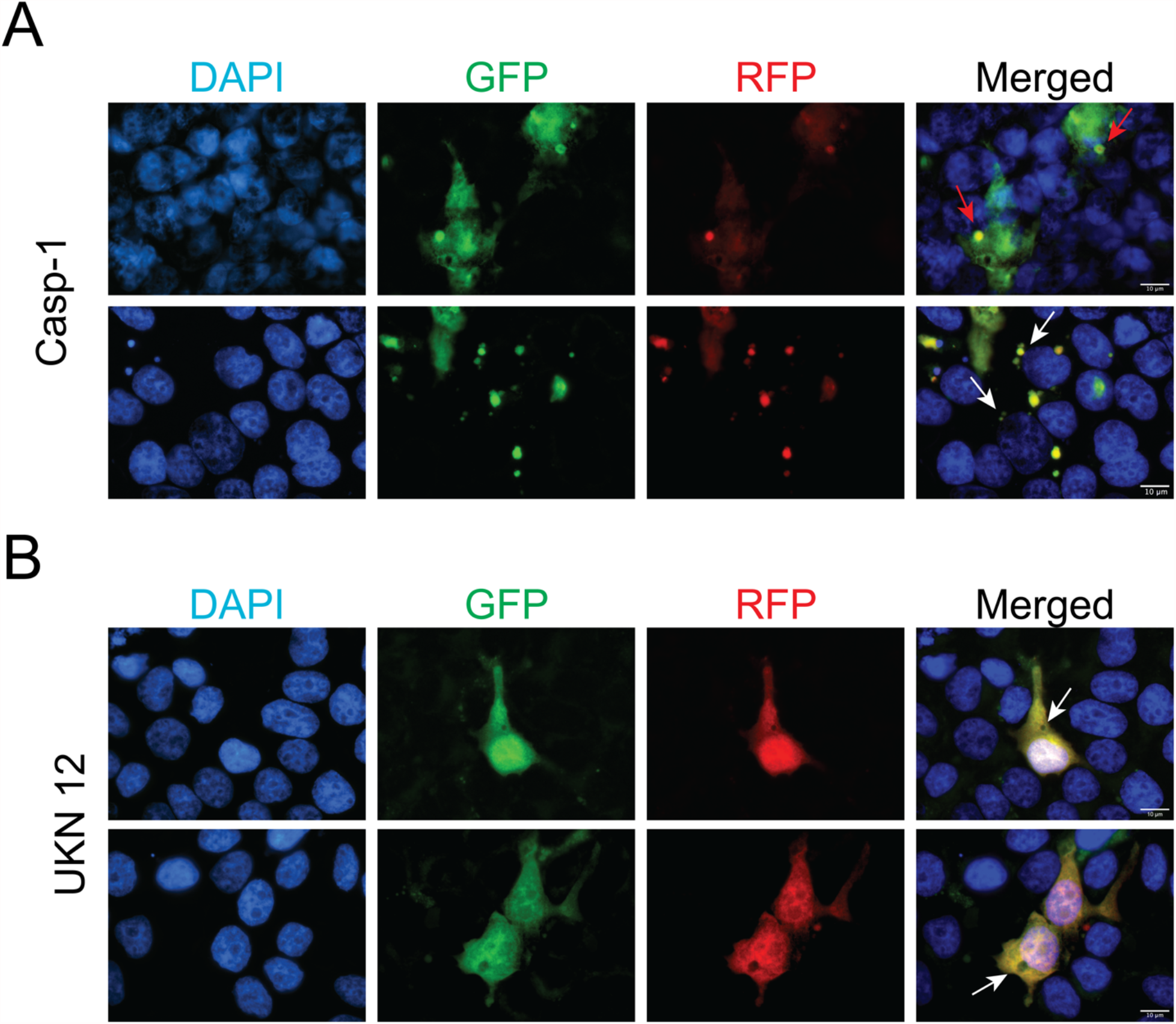
Bi-fluorescent reporter recapitulates FLICA-like staining of active caspase-1. HEK293T cells were transfected with 100 ng myc-ASC, 100 ng NLRP3, 400 ng p-Casp1 (WT or mutant) and 400 ng GFP-YVAD-RFP. 18 hrs post-transfection, cells were fixed analyzed for GFP-RFP aggregates by microscopy. **A**. Micrographs for GFP (Green), RFP (Red) and Merged with DAPI (Blue) showing distribution of active caspase-1 sites in a cell. (Top) The bi-fluorescent reporter showing speck-like morphology (red arrows). (Bottom) The bi-fluorescent reporter showing multiple GFP-RFP aggregates (White arrows). **B**. Micrographs showing a reporter-free region close to nucleus (white arrow; expected site of the speck). Representative images from three independent experiments.

**Figure S5:**
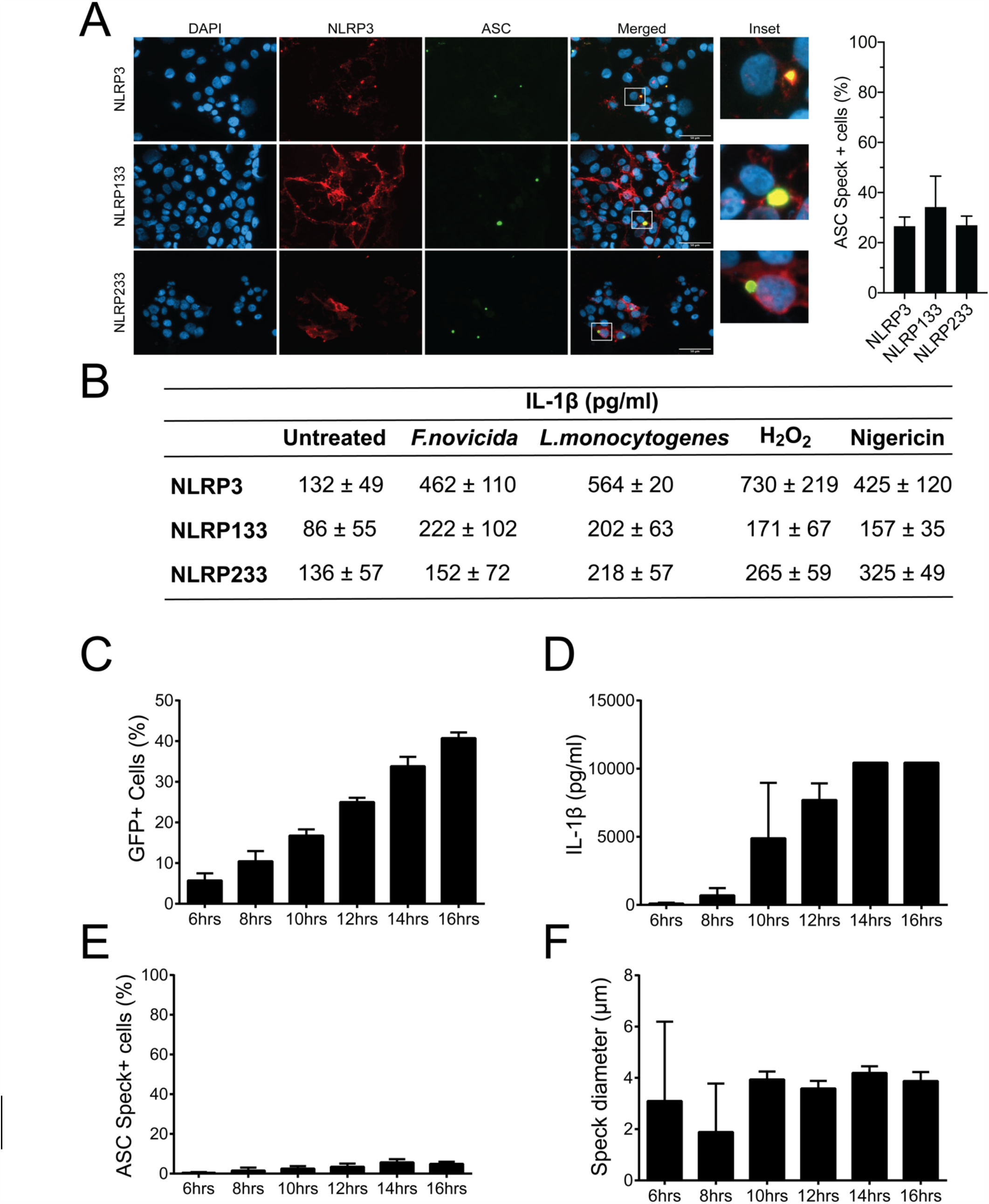
Uncoupling of speck formation and IL-1β processing. **A**. HEK293T cells were transfected with 1 µg myc-ASC and 1µg Flag-NLRP3 or a chimeric Flag-NLRP133 or Flag-NLRP233. After 18 hrs cells, fixed and permeabilized cells were stained with anti-Flag (M2) and anti-ASC (Santa Cruz) and then analyzed for speck formation. Percentage of cells containing specks. Minimum of 140 cells were counted from 4 individual field of view; for NLRP233, 30 cells were counted. Data is presented as Mean ± SEM and analyzed using one-way ANOVA followed by Tukey’s multiple comparison test. **B**. IL-1β processing in inflammasome reconstituted HEK293Ts expressing NLRP3 or the indicated chimeras and treated/infected with stated NLRP3 agonists (See **Methods**). **(C-F)** HEK293T cells were transfected with 100 ng NLRP3, 100 ng GFP-ASC, 20 ng p-Caspase-1 and 200 ng pro-IL-1β allowing agonist-independent maturation and release of active IL-1β. At stipulated time-points, cells and culture supernatants were harvested. Cells were fixed and analyzed for GFP expression **(C)**, Culture supernatant was analyzed for released IL-1β **(D)**, ASC speck **(E)** and speck diameter **(F)**.

## ACKNOWLEDGEMENTS

We thank Drs. Jim Drake and William O’Connor (Albany Medical College, Albany, NY) for their insightful comments on initial drafts of the manuscript. Adobe Illustrator 2020 and Servier Medical Art templates have been used to draw the schematics.

